# Three-dimensional electron microscopy uncovers mitochondrial network remodeling in residual triple negative breast cancer persisting after conventional chemotherapy treatments

**DOI:** 10.1101/2024.09.09.611245

**Authors:** Mariah J. Berner, Heather K. Beasley, Benjamin Rodriguez, Audra Lane, Steven W. Wall, Andrea G. Marshall, Zer Vue, Larry Vang, Mokryun L. Baek, Mason Killion, Faben Zeleke, Bryanna Shao, Dominque Parker, Autumn Peterson, Julie Sterling Rhoades, Lacey E. Dobrolecki, Amber Crabtree, Annet Kirabo, Sepiso Mansenga, Prasanna Katti, Prasanna Venkhatesh, C.V. Dash, Michael T. Lewis, Antentor Hinton, Gloria V. Echeverria

**Affiliations:** Lester and Sue Smith Breast Cancer, Baylor College of Medicine, Houston, TX, USA; Dan L Duncan Comprehensive Cancer Center, Baylor College of Medicine, Houston, TX, USA; Department of Medicine, Baylor College of Medicine, Houston, TX, USA; Department of Molecular Physiology and Biophysics, Vanderbilt University, Nashville, TN, USA; Department of Veterans Affairs, Tennessee Valley Healthcare System, Nashville, TN, USA; Program in Cancer Biology, Vanderbilt University School of Medicine, Nashville, TN, USA; Department of Medicine, Division of Clinical Pharmacology, Vanderbilt University Medical Center, Nashville, TN, USA; First Center for Autism & Innovation, Vanderbilt University, Nashville, TN, USA; Department of Biology, Indian Institute of Science Education and Research (IISER), Tirupati, AP, India; The Center for AIDS Health Disparities Research, Meharry Medical College, Nashville, TN, USA; Department of Microbiology, Immunology, and Physiology, Meharry Medical College, Nashville, TN, USA; Department of Biochemistry, Cancer Biology, Pharmacology and Neuroscience, Meharry Medical College, Nashville, TN, USA; Department of Molecular and Cellular Biology, Baylor College of Medicine, Houston, TX, USA; Department of Radiology, Baylor College of Medicine, Houston, TX, USA; Department of Medicine, Division of Cardiology, Vanderbilt University Medical Center, Nashville, TN, USA; Department of Radiation Oncology, Baylor College of Medicine, Houston, TX, USA

## Abstract

Mitochondria are hubs of metabolism and signaling, playing crucial roles in tumorigenesis, therapeutic resistance, and metastasis in many types of cancer. Various laboratory models of cancer demonstrate the extraordinary dynamics of mitochondrial structure, but little is known about the full extent of the complexity of the mitochondrial network, nor its regulation upon exposure to therapeutic stressors. We previously demonstrated the importance of mitochondrial structure and oxidative phosphorylation in the survival of chemotherapy-refractory triple negative breast cancer (TNBC) cells. As TNBC is a highly aggressive breast cancer subtype with few targeted therapy options, conventional chemotherapies remain the backbone of TNBC treatment. Unfortunately, approximately 45% of TNBC patients retain substantial residual tumor burden following chemotherapy, associated with abysmal prognoses. Herein we present the first three-dimensional analysis of mitochondrial networks in human tumor tissues. Use of two experimentally tractable orthotopic patient-derived xenograft (PDX) models TNBC enabled us to conduct longitudinal analyses to construct mitochondrial networks in treatment-naïve and residual tumors persisting after exposure to a variety of conventional chemotherapies. Further, we modeled lipid droplet (LD) structures and their physical contacts with mitochondria. In total, we reconstructed 3,750 mitochondria and 800 LDs in three dimensions using serial block-face scanning electron microscopy (SBF-SEM), providing unprecedented insights into the complexity and intra-tumoral heterogeneity of mitochondria in TNBC. Both carboplatin (CRB) and docetaxel (DTX) chemotherapies produced residual tumors that harbored mitochondria with significantly increased areas, volumes, and perimeters in both PDX models. Additionally, treatment with the conventional combinations DTX plus CRB or Adriamycin plus cyclophosphamide (AC), led to reduced mitochondrial branching and elongation. In contrast, DTX or CRB alone elicited model-specific changes in mitochondrial complexity. Further, the extensive intra-tumoral heterogeneity of mitochondrial structure in untreated PDX tumors significantly decreased in residual tumors. Analyses of LDs revealed significant and consistent elevation of the number and physical proximity of MLCs in residual tumors, congruent with our previous studies providing evidence for transcriptomic and proteomic rewiring of lipid metabolism in residual TNBC. These results highlight the potential for structure-based monitoring of chemotherapeutic metabolic reprogramming and suggest unique molecular mechanisms that may underlie chemoresistance in TNBC. Furthermore, our findings provide novel insights into a new type of intratumoral heterogeneity, that of mitochondrial intratumoral heterogeneity, which complements our understanding of the genomic, epigenomic, transcriptomic, and proteomic complexity of TNBC.

## INTRODUCTION

Triple negative breast cancer (TNBC), an aggressive subtype of breast cancer, comprises approximately 15% of breast cancer cases^1^. Because TNBCs do not express estrogen or progesterone receptors or overexpress the human epidermal growth factor receptor 2, there are limited targeted therapies for this subtype^1,2^. Most patients with TNBC receive pre-surgical neoadjuvant chemotherapy (NACT) typically including several conventional chemotherapeutic agents given in simultaneous and/or sequential combinations. NACT now also consists of the immunotherapy agent Programmed Cell Death 1 (PD-1) inhibitor, with a marginal but statistically significant improvement in response rate^3^. Conventional chemotherapies typically include DNA-damaging agents (*e.g*., Adriamycin, cyclophosphamide, carboplatin) and microtubule disrupting taxanes (*e.g.,* docetaxel, paclitaxel). As there is no single accepted standard NACT regimen for TNBC, a variety of regimens are used throughout the United States and worldwide^4^. Pathology evaluation of the resected surgical specimen assesses residual cancer burden (RCB) following NACT. Unfortunately, approximately 45% of TNBC patients retain substantial RCB after NACT, which is strongly associated with high rates of metastatic relapse and mortality^5,6^. Thus, understanding the molecular features of residual tumor cells that survive chemotherapeutic regimens is key to developing novel strategies to combat residual disease.

Mitochondria adapt to meet the cell’s energetic and signaling demands in response to exogenous stressors, in part through mitochondrial structural rearrangements and communication with other organelles^7^. In the context of cancer, metabolic adaptations orchestrated by mitochondria can encompass oxidative phosphorylation (oxphos), glycolysis, reactive oxygen species (ROS), nucleotide synthesis, and fatty acid oxidation as we recently reviewed^8^. The interplay between mitochondrial fission and fusion, dictating mitochondrial structure, has been studied in relation to cancer therapy resistance and metastasis. For TNBC, mitochondrial structure dynamics play a role in metastatic phenotypes, although the findings seem to be context-dependent^9,10^, as well as resistance to standard chemotherapies^11^. Importantly, we demonstrated that chemotherapy-induced adaptations in mitochondrial structure and function produced a new therapeutic dependency for residual TNBC, allowing us to delay tumor regrowth by inhibiting mitochondrial fusion or by direct inhibition of Complex I^11–13^. Complementing our TNBC work^11^ with the first-generation OPA1 inhibitor MYLS22^14^, an optimized compound was recently described^15^, further establishing the promise of inhibiting mitochondrial structure dynamics as ananticancer strategy. These studies highlight the growing interest in translating mitochondrial structure perturbations to the clinic for cancer patients.

Serial block-face scanning electron microscopy (SBF-SEM) was first developed in 2004 by Denk and Horstmann to study mitochondrial structure and location within the neuron^16,17^. This state of the art imaging method produces high-resolution micrographs that capture subcellular ultrastructural features through three-dimensional (3D) rendering that, unlike other volumetric electron microscopy techniques, provides large x- and y-axis ranges^18–20^. These visualizations of changes in mitochondrial morphology, orientation, configuration, size, and shape provide insights into mitochondrial physiology and function, important for physiology and disease^21–24^. We and others have shown that using 2D transmission electron microscopy (TEM) to characterize mitochondrial structures in breast cancer provides useful, albeit limited, insights into the extent of the mitochondrial network^10,11,25,26^. Previous SBF-SEM studies of mitochondria demonstrated dynamic 3D structures and networks associated with skeletal muscle physiology^27^, aging^28–30^, and tumor cell culture methodologies^31^. Of note, a recent non-small cell lung cancer study employed genetically engineered mouse models to demonstrate that 3D mitochondrial structure was spatially heterogeneous within tumor cells and functionally affected mitochondrial respiration^32^.

The nature of 3D mitochondrial networks in human cancer tissues remains currently undocumented. In fact, there are no published SBF-SEM mitochondria analyses of any human cancer tissues. Further, this approach has not been used to address biologically important transitions during cancer, namely response to therapeutic stressors. To address these knowledge gaps in light of our prior work demonstrating the importance of mitochondrial structure and oxphos for TNBC chemoresistance, we used SBF-SEM to determine 3D mitochondrial networks in two experimentally tractable orthotopic PDX models of TNBC. PDXs provide valuable models for longitudinal studies to evaluate chemotherapy response as they effectively recapitulate the molecular features and heterogeneity observed in human TNBC^33–37^. Our longitudinal analyses enabled us to compare residual tumors surviving exposure to a variety of conventional chemotherapies to their treatment-naïve counterparts. We found significant changes in mitochondrial shape, size, branching, complexity, and contacts with lipid droplets (LDs) in residual tumors compared to treatment-naïve tumors. Further, we found striking intratumoral heterogeneity (ITH) of mitochondria (mITH) within individual tumors that was most pronounced in the treatment-naïve tumor. This finding highlights the incredible complexity within individual TNBC tumors and reaffirms the extensive ITH that has been thoroughly documented at the genomic, transcriptomic, epigenetic, and proteomic levels^38,39^. Together, these results demonstrate the incredible adaptability of mitochondrial 3D structure as TNBCs are exposed to conventional chemotherapies. We and others have shown the promise of mitochondrial interventions for anticancer therapy, and these results lay the groundwork for structure-based monitoring of treatment responses and anti-mitochondria therapeutic efficacy in TNBC.

## METHODS

### Patient-derived xenograft mouse studies

Animal studies were conducted in accordance with the National Institutes of Health *Guide for the Care and Use of Laboratory Animals* with approval of the BCM IACUC under protocol AN-8243 and AN-2289. Mice were ethically euthanized as recommended by the Association for Assessment and Accreditation of Laboratory Animal Care. The PIM001-P PDX model was originally obtained from the University of Texas MD Anderson Cancer Center, where it was generated and characterized and was then propagated at BCM as described previously^13,35^. Briefly, cryo-preserved PDX cells were suspended as a 1:1 volume mixture in a total of 20 µl of complete cell growth medium (Dulbecco’s modified Eagle’s medium:F12, Cytiva HyClone, SH30023.01) supplemented with 5% fetal bovine serum and Matrigel (Corning, 354234). 5.0 × 10^5^ viable tumor cells were injected unilaterally into the fourth mammary fat pads of 5- to 8-week-old female Nod-Rag-Gamma mice (NRG*, Nod.Cg-Rag1* ^tm1Mom^ */IL2rg* ^tm1wjl^ */SzJ*, The Jackson Laboratory). The WHIM14 PDX model mice were implanted unilaterally as 1×1 tumor chunks into the fourth mammary fat pad by the BCM Patient-derived Xenograft Core as described previously^40^.

To validate PDX models, the Cytogenetics and Cell Authentication Core at MD Anderson Cancer Center performed short-tandem repeat (STR) DNA fingerprinting on DNA extracted from PDX tumors. The Promega 16 High Sensitivity STR Kit (Catalog # DC2100) was used for fingerprinting analysis, and profiles were compared to online databases (DSMZ/ ATCC/ JCRB/ RIKEN).

Mouse body weights and tumor volumes, measured using digital calipers, were recorded twice weekly for the duration of the experiment. When mammary tumors reached an average size of 150 mm^3^, mice were randomized into five treatment groups (n = 6 mice per group) to monitor tumor responses and collect samples for molecular analyses. For the treatment-naïve group, the tumors were harvested at 150 mm^3^ without any treatment. Mice in the Adriamycin plus cyclophosphamide (AC) treatment group received one dose of Adriamycin and cyclophosphamide every three weeks for a total of three doses. Adriamycin (doxorubicin, ChemieTek, CT-DOXO) was solubilized in sterile water immediately before administration and protected from light prior to intraperitoneal (i.p.) injection at a dose of 0.5 mg/kg. Cyclophosphamide (Sandoz, 0781-3244-94) was solubilized in sterile water immediately prior to administration via i.p. injection at 50 mg/kg. Mice in the docetaxel (DTX), the carboplatin (CRB), and the docetaxel plus carboplatin combination (DTX+CRB) treatment groups received weekly doses for three weeks. Docetaxel (Hospira, 0409-0201-10) was supplied in a solution of polysorbate 80NF, 4 mg anhydrous citric acid USP, 23% v/v dehydrated alcohol USP, q.s. with polyethylene glycol 300 NF at 20 mg/kg. DTX was administered by i.p. injection at a dose volume of 2 ml/kg. Carboplatin (TEVA Pharmaceuticals, 0703-4248-01) was supplied at 10 mg/ml in water for i.p. injection at 50 mg/kg at a dose volume of 5 ml/kg. For the DTX+CRB treatment group, each compound was administered consecutively by i.p. injection. For each group of the PIM001-P PDX model, residual tumors were harvested one complete cycle after the third dose of chemotherapy (i.e., a total of 28 days after the start of treatment for DTX, CRB, DTX+CRB, and 63 days after the start of treatment for AC). For each group of the WHIM14 PDX model, residual tumors were harvested when they had regressed in size by 40% because insufficient tumor material would be available for analyses after ∼1.5 weeks of starting treatment in this model due to its rapid regression.

### Formalin-fixed paraffin-embedded (FFPE) tissue staining

After harvesting the tumors, we immediately fixed tumor fragments in formalin (Sigma HT501128-4L) for 48 hours at room temperature on a rocker, washed them with 1× phosphate-buffered saline, and stored them in 70% ethanol at 4 °C until they were submitted to the BCM Pathology Core and Lab to be processed into paraffin blocks, cut into 3um sections, and stained with hematoxylin and eosin (H&E). Immunohistochemical staining with antibodies against Ki67 (Dako catalog #M7240, 1:200), Ku80 (CST 2180, 1:100), and human-specific mitochondria (Abcam 92824, 1:1,000) was conducted as described previously^13^.

### SBF-SEM sample processing

Tumors were harvested from the mammary gland immediately after the mice were euthanized and immediately placed in fixative. A tumor slice ∼1 mm thick was placed into 1 ml of fixative (2% paraformaldehyde + 2% glutaraldehyde in 0.15 M cacodylate buffer) and stored at 4°C until they were sent to the Mayo Microscopy and Cell Analysis Core for processing and imaging. Tissue samples for SBF-SEM were prepared using a modified procedure^23,41^. Briefly, fixed tissue was rinsed in 0.1 M cacodylate buffer and placed in 2% osmium tetroxide + 1.5% potassium ferracyanide in 0.1 M cacodylate, washed with nH_2_O (MilliQ water), incubated at 50°C in 1% thiocarbohydrazide, incubated again in 2% osmium tetroxide in nH_2_O, rinsed in nH_2_O and placed in 2% uranyl acetate overnight. The next day, the tissue was rinsed again in nH_2_O, incubated with Walton’s lead aspartate, dehydrated through an ethanol series, and embedded in Embed 812 resin. To prepare embedded tissue for placement into the SBF-SEM, a ∼1.0 mm^3^ piece was trimmed of excess resin and mounted on an 8 mm aluminum stub using silver epoxy Epo-Tek (EMS, Hatfield, PA). The mounted sample was then carefully trimmed into a smaller ∼0.5 mm^3^ tower using a Diatome diamond trimming tool (EMS, Hatfield, PA) and vacuum sputter-coated with gold-palladium to help dissipate any charge. Sectioning and imaging of the sample were performed using a VolumeScope 2 SEM^TM^ (Thermo Fisher Scientific, Waltham, MA). Imaging was performed under low vacuum/water vapor conditions with a starting energy of 3.0 keV and a beam current of 0.10 nA. Sections of 50 nm thickness were cut, providing a final image spatial resolution of either 15 nm × 15 nm × 50 nm or 10 nm × 10 nm × 50 nm.

### SBF-SEM data analysis

Image analyses, including registration, volume rendering, and visualization, were performed using Amira (Thermo Fisher) software packages. The selection criteria for a region of interest (ROI) included tumor regions and excluded necrotic and extracellular matrix regions. A minimum of three ROIs were selected per sample. For each sample, images were collected from at least 300 serial sections that were then stacked, aligned, and visualized using Amira to make videos and quantify volumetric structures. A random number generator was used to select 350 segmented mitochondria for each tumor of the PIM001-P for the final analysis. See Supplementary Table 1 for the specific number of mitochondria segmented per ROI and those selected for final analysis of the PIM001-P model. With the WHIM14 model, two ROIs were selected per sample, and a random number generator was used to select 500 segmented mitochondria for each WHIM14 tumor (supplementary Table 2). Additionally, for the WHIM14 model, we segmented a total of 200 lipid droplets from the two selected ROIs to analyze the mitochondria-lipid droplet interactions further (supplementary Table 3). Previously defined protocols using Amira software provide a workflow for 3D reconstructed organelles by manual or semi-automated segmentation^19,20^. Amira provides parameters for quantifying 3D structures, many of which are included in the software, such as volume and surface area. We have added sphericity, which is measured through the Feret’s Diameter. Feret’s diameter refers to the distance between any two most distant points in a 3D object. Sphericity refers to the adherence of the object to that of a sphere and is calculated by the equation (π^1/3^(6*V*)^2/3^)/*SA*, where *V* is the volume of the object, and *SA* is the surface area of the object. We also quantified the mitochondrial complexity index, calculated by the equation *SA*^3^/16π^2^*V*^2^, and the mitochondrial branching index, which calculates the relative branching between the transverse and longitudinal mitochondrial orientations^24^.

### Statistical analyses

We evaluated the means for each measurement between treatment groups by the Mann–Whitney U test followed by the Holm method of p-value adjustment^42–44^. We performed Bartlett’s test followed by pairwise F-tests to determine whether the variance of each measurement differed between treatment groups using R 4.1.0 (R Core Team, 2023) and the *stats* package (base statistical package in R)^45^. Bartlett’s test assesses whether all the treatment groups have the same variance. A p-value less than 0.05 for a given measurement by Bartlett’s test indicates that some of the treatment groups have different variances for that measurement. To identify which treatment groups differed, we then conducted a two-tailed F-test on each possible pairwise comparison between treatment groups. In a two-tailed F test, the F value is defined as the quotient of the variance of one group divided by the variance of the other. This F value is used to test whether the variance in the numerator equals that in the denominator. For a given F test, a p-value was determined by taking the area under the F distribution beyond that test’s F value and then subtracting that area from 1. Once all possible comparisons had been made for a measurement, the p-values for that measurement were adjusted by the Holm method to account for multiple comparisons.

## RESULTS

### Various conventional chemotherapies elicited drastic tumor regression followed by regrowth in TNBC PDX models

To model the residual tumor state, which we previously showed exhibits increased OXPHOS and mitochondrial dependence, we treated the orthotopic PIM001-P^13,35^ PDX model with conventional chemotherapy agents administered either individually or in combination: AC, docetaxel (DTX), carboplatin (CRB), or DTX+CRB (Fig. 1a). Interestingly, we found that DTX, CRB, and DTX+CRB all elicited similar responses as did AC, congruent with our previous reports: initially drastic tumor regression to a ‘residual’ tumor, maintenance of a residual tumor for weeks to months, then regrowth of the tumor. We next tested an additional orthotopic PDX model, WHIM14^46^, with DTX, CRB, or DTX+CRB, similarly demonstrating tumor regression followed by regrowth (Fig. S1a). Both PIM001-P and WHIM14 are models of aggressive TNBC: PIM001-P was derived from the primary tumor of a treatment-naïve TNBC patient diagnosed with *de novo* metastasis, who received gemcitabine and paclitaxel had an overall survival of four months^13,35^. WHIM14 was generated from a chest wall metastasis of a heavily pre-treated TNBC patient who continued to receive additional therapies, with an overall survival of 37 months^46^.

**Figure 1.**
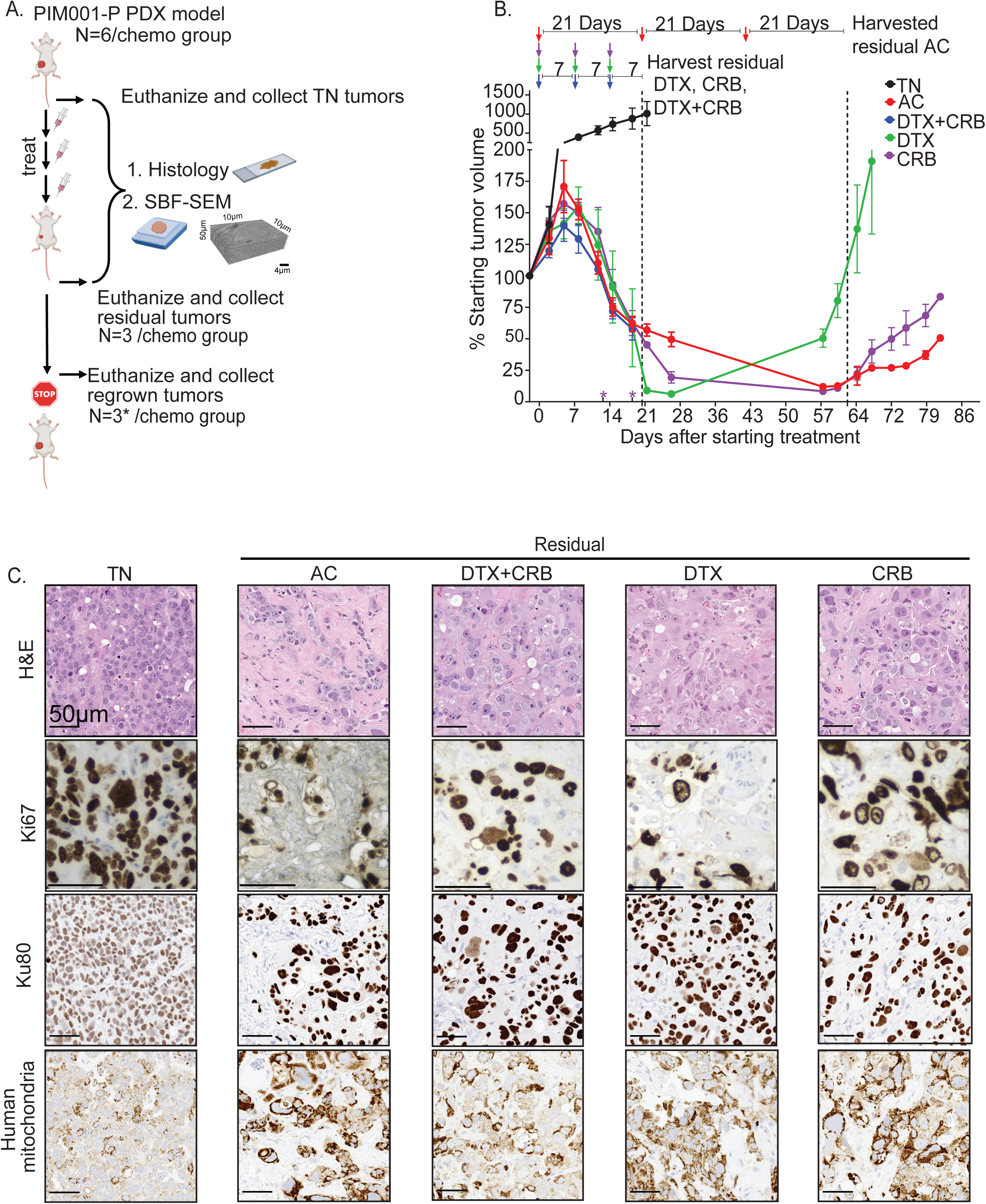
PIM001-P PDX tumors persisted following conventional chemotherapy treatments. (a) Schematic of the experimental approach used in the study. Images were adapted from BioRender. (b) Tumor volumes were monitored biweekly following administration of chemotherapies to mice bearing orthotopic PIM001-P tumors (n = 6 mice/group). Treatment groups were treatment-naïve (TN), Adriamycin + cyclophosphamide (AC), docetaxel + carboplatin (DTX+CRB), docetaxel (DTX), and carboplatin (CRB). Arrows at the top indicate when chemotherapy was administered. Asterisks (*) above the x-axis indicate early euthanasia due to animal health concerns. Residual tumors were harvested, as noted by arrows. Error bars represent the standard error of the mean. (c) Residual tumors were harvested, processed for FFPE, and analyzed by H&E staining and IHC for Ki67, Ku80 and human-specific mitochondria. Scale bars are 50 µm.

Treatment-naïve (TN) tumors were collected from a subset of mice for each model when tumors reached approximately 150 mm^3^. For PIM001-P, after three treatment cycles, residual tumors were harvested on the day that the fourth cycle would have begun. These residual tumors showed an average decrease in tumor volume of 53.6% (DTX, 52.9%; CRB, 61.0%; DTX+CRB, 61.5%; and AC, 34.0%) (Fig. 1b). WHIM14 regressed more rapidly and extensively, with an average reduction in tumor volume of 60% after only one cycle of treatment (Fig. S1a). Accordingly, ‘residual’ tumors in this model were harvested relatively early, when volumes had decreased to approximately 40% of the initial volume after the first cycle. The mice remaining in each treatment arm for regrowth monitoring received three additional cycles (four in total) of chemotherapy. At the indicated time points, three to four mice were euthanized, and their residual tumors were collected, and from these, one tumor was randomly selected for SBF-SEM and conventional histology analyses. The remaining mice were monitored for relapse and then euthanized once tumors regrew to at least 100% of their original volume. In three cases, mice succumbed to drug-related toxicities, all within the DTX+CRB group. We were unable to assess the regrowth in PIM001-P after four cycles of DTX+CRB. To address this, we conducted a follow-up study with a single cycle of treatment, demonstrating robust regrowth after the initial regression (Fig. S2a).

H&E staining of TN FFPE tumor sections revealed poorly differentiated cellular architecture with abundant mitotic figures. All residual tumors in chemotherapy treatment groups exhibited extensive desmoplasia with significant tumor cell pleomorphism, including atypical and enlarged nuclear and cytoplasmic sizes and shapes, pronounced nucleoli, and areas of necrosis, consistent with chemotherapy-induced cytological changes observed previously in PDXs and in patients^13,47^ (Fig 1c & S1b). Notably, residual tumors retained cycling Ki67 positive cells (Fig 1c), in line with our prior AC study in PIM001-P^13^. Ku80 staining, used to distinguish human tumor cells from mouse stromal cells, revealed an abundant stromal component in all residual tumors (Fig. 1c), as previously reported with AC treatment^13^. Staining with an antibody specific for human mitochondria, demonstrated increased mitochondrial content in residual tumor cells compared to TN cells, consistent with our prior finding in AC-treated residual tumors (Fig. 1c)^13^. In summary, various conventional chemotherapies induced tumor regression in both PIM001-P and WHIM14. The residual tumors exhibited regrowth capacity, aberrant histology architecture, and increased mitochondrial content in the tumor cell compartment.

### 3D rendering of mitochondria by SBF-SEM revealed residual tumor mitochondria had increased volume, area, and perimeter relative to TN

From each treatment group, we selected one representative tumor with a volume close to the group’s mean to generate blocks for SBF-SEM. Electron micrographs were acquired at either 15 nm × 15 nm × 50 nm or 10 nm × 10 nm × 50 nm resolution with a minimum of three ROIs, each with 300 slices, per tumor. ROIs were inspected prior to imaging to avoid necrotic and stroma-rich regions. Mitochondria segmentation was conducted across different areas of each orthogonal (ortho) slice (Fig. 2a-e; Fig. S3a-d) to capture spatial heterogeneity within tumors. Each individual segmented mitochondrion used for final analyses was manually inspected to confirm complete segmentation (*i.e.,* not missing a slice) (Fig. 2a’-e’, Fig. S3a’-d’). From each tumor, we randomly selected 350 mitochondria from PIM001-P (Fig. 2a’-e’’) and 500 mitochondria from WHIM14 (Fig. S3a’’-d’’) for morphometric analyses from a minimum of three or two ROIs, respectively. For each mitochondrion, we computed area, volume, and perimeter as described previously^19,20,27^.

**Figure 2.**
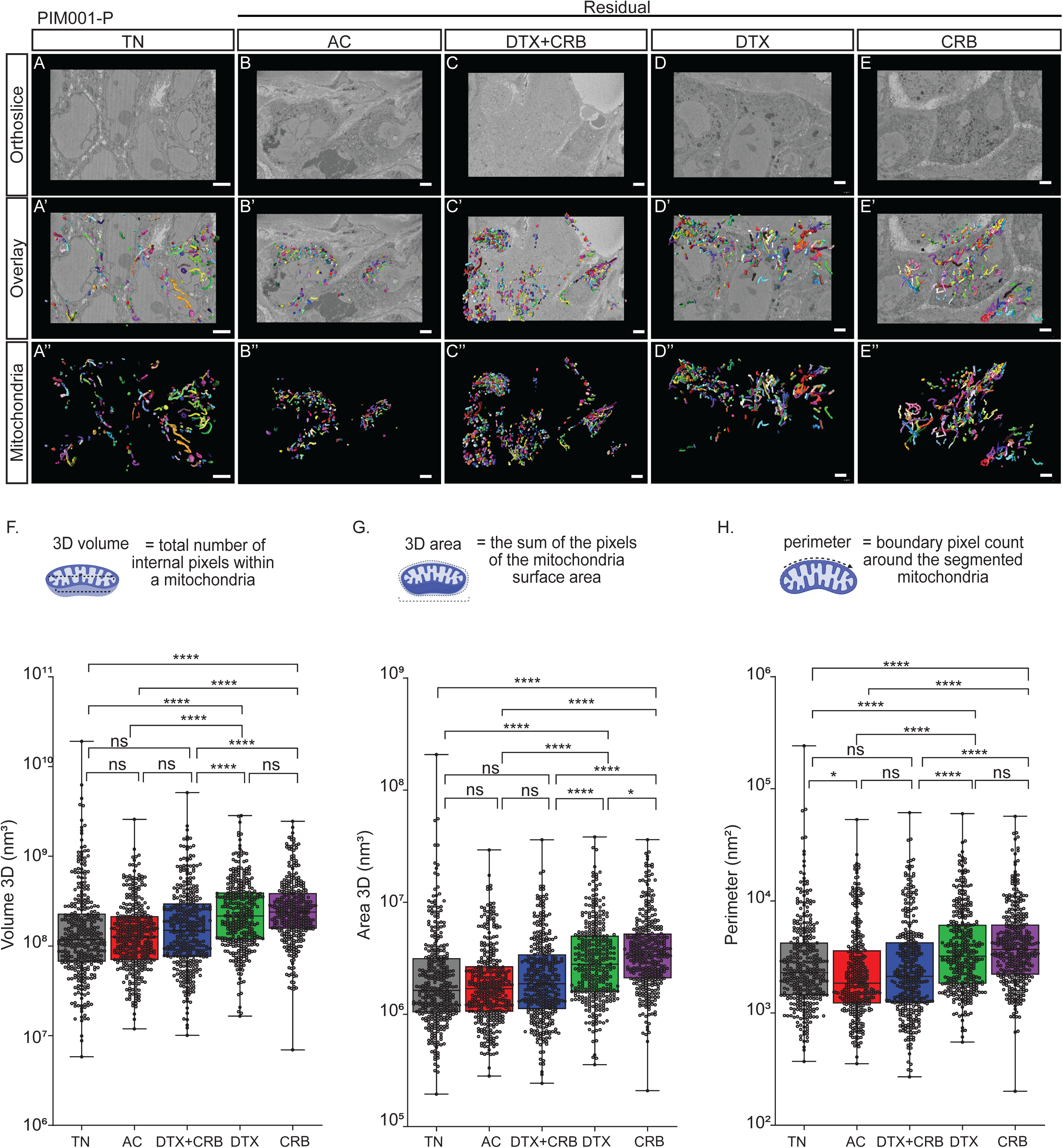
Single-agent chemotherapy treatments in PIM001-P significantly increased mitochondrial size in residual tumors relative to TN. (a-e) Representative SBF-SM orthoslices. (a’-e’) Representative 3D SBF-SEM orthoslices with overlays of mitochondria segmentations. (a’’-e’’) Representative 3D SBF-SEM mitochondria segmentations for each treatment group: treatment-naïve (TN), Adriamycin + cyclophosphamide (AC), docetaxel + carboplatin (DTX+CRB), docetaxel (DTX), and carboplatin (CRB). The scale bar is 3 µm. Mitochondrial measurements were calculated using Amira software for 350 segmented mitochondria (represented as dots) from each tumor for (f) 3D volume, (g) 3D area, (h), and perimeter. Adjusted p-values (*p < 0.05, **p < 0.01, ***p < 0.001, ****p<0.0001) are shown on box and whisker plots as determined using the Mann–Whitney U test followed by the Holm method.

We compared the mean values of each mitochondrial feature between tumors. To assess differences in these means between treatment groups, we performed Mann–Whitney U tests, as q–q plots indicated that mitochondrial features were not normally distributed, precluding the use of ANOVA (Fig. S4). In both PDX models, DTX and CRB residual tumors had significantly increased mitochondrial area, volume, and perimeter compared to treatment-naïve tumors (Fig 2f-h, S3e-g). This effect was also observed following DTX+CRB combination treatment in WHIM14. Although no significant changes were detected in PIM001-P with the DTX+CRB combination, the median mitochondrial area and volumes were elevated relative to TN. In contrast, AC treated PIM001-P residual tumors exhibited no change in mitochondrial area, perimeter, or volume relative to TN.

TNBC is characterized by extensive intratumoral heterogeneity at the genomic^48–50^, epigenetic^51^, transcriptomic^52^, cell type^53^, spatial^54,55^, and proteomic^56,57^ levels. Here we observed substantial intra-tumoral mitochondrial heterogeneity (mITH), which was more pronounced in PIM001-P than in WHIM14. To quantify this, we generated density plots and computed variances of each metric in each sample. The density plots confirmed the aforementioned increases in area, volume, and perimeter and further illustrated a broader distribution of mitochondrial features in TN tumors compared to most residual tumors (Fig. S5-8). Notably, variance was significantly reduced in all residual tumors compared to TN in both models, with the exception of mitochondrial volume in DTX-treated WHIM14 (Fig S9 & S10). This reduction in variance was particularly striking in PIM001-P (Fig S9). Together, these findings demonstrate that DTX, CRB, and DTX+CRB elevated mitochondrial area, volume, and perimeter in both PDX models while simultaneously inducing a significant reduction in mITH.

### Residual tumors harbored mitochondria with greater complexity

We next analyzed mitochondrial elongation and branching. Sphericity scores (ranging from 0 to 1, with 1 representing a perfect sphere) were calculated, revealing significant yet minor changes in residual tumors relative to TN tumors. (Fig. 3a&b, Fig. S11a&b). Specifically, mitochondria in all residual WHIM14 tumors, as well as those in the DTX+CRB and AC-treated residual PIM001-P tumors, showed increased sphericity (significantly, or trending towards significance). By contrast, mitochondria in PIM001-P residual tumors following DTX and CRB treatment had significantly decreased sphericity, consistent with elongation. To further evaluate elongation, we computed the mitochondrial branching index (MBI), a conventional metric that reflects the directionality of elongation^27^. MBI is defined as the ratio of the longest point in the transverse direction (in this case, width) to the longest point in the longitudinal direction (length) (Fig. S12a). MBI results matched the aforementioned sphericity results (Fig. S12b&c). To assess mitochondrial branching and structural complexity, we calculated the mitochondrial complexity index (MCI)^27^, which incorporates both 3D volume and surface area. Conceptually, a more branched mitochondrion would have a larger surface area than a less branched one of equal volume, yielding a higher MCI. Residual tumor mitochondria in WHIM14, as well as those in the DTX+CRB and AC-treated PIM001-P residual tumors, showed significant decreases in MCI. In contrast, mitochondria from PIM001-P tumors treated with single agent DTX or CRB treatment increased MCI (Fig. 3c & S11c), consistent with greater branching and/or elongation.

**Figure 3.**
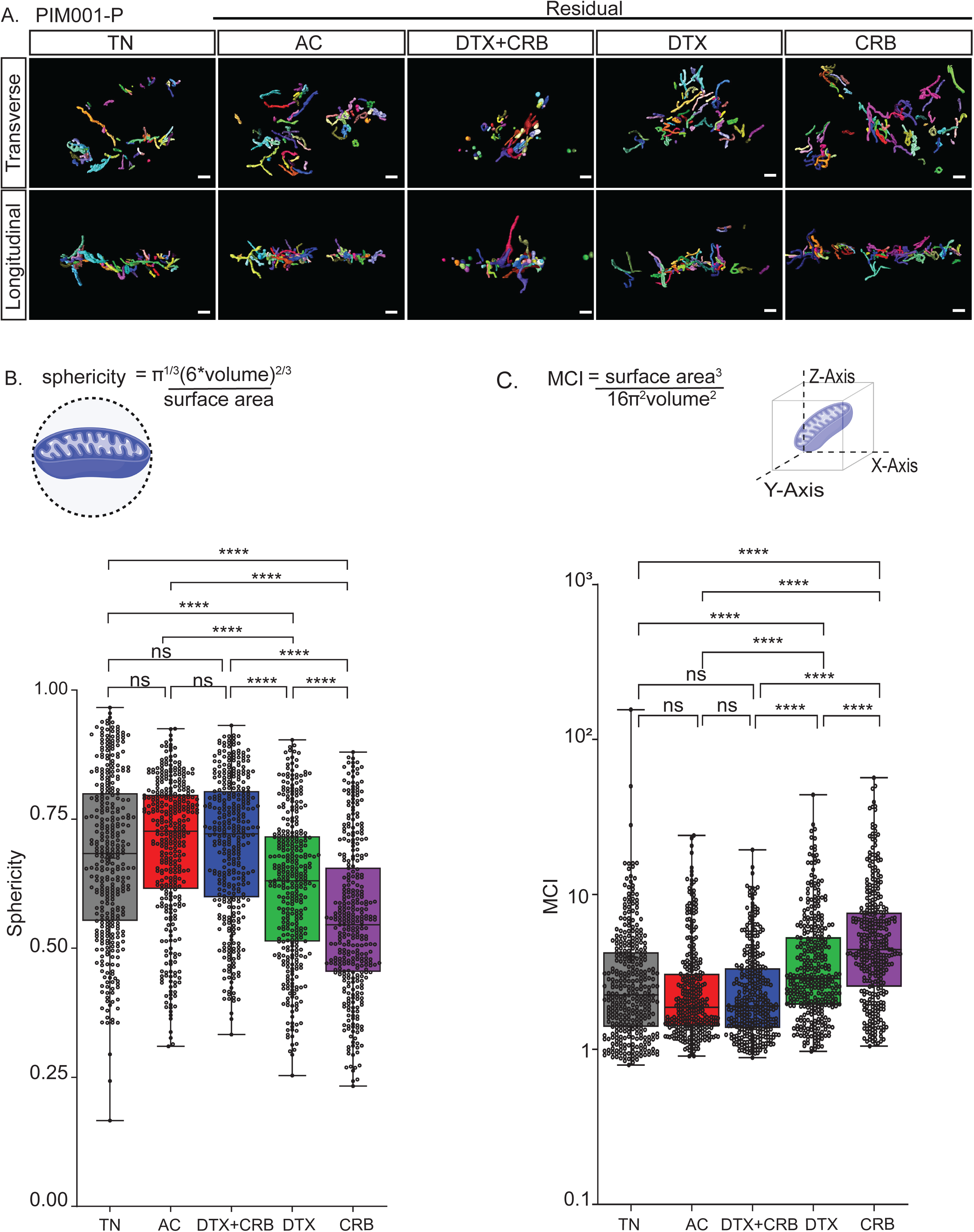
Single-agent chemotherapy treated residual tumors’ mitochondria are more branched and less spherical relative to TN in PIM001-P. Mitochondrial networks revealed through 3D mitochondria rendered using SBF-SEM. (a) Representative longitudinal and transverse views of the segmented mitochondria for each treatment group: treatment-naïve (TN), Adriamycin + cyclophosphamide (AC), docetaxel + carboplatin (DTX+CRB), docetaxel (DTX), and carboplatin (CRB). The scale bar is 3 µm. Mitochondrial measurements were calculated using Amira software for 350 segmented mitochondria (represented as dots) from each tumor for (b) sphericity (c) mitochondrial complex index (MCI), along with their respective method of measurements below. Adjusted p-values (*p < 0.05, **p < 0.01, ***p < 0.001, ****p<0.0001) are shown on box and whisker plots as determined using the Mann–Whitney U test followed by the Holm method.

We next assessed mitochondrial shape using MCI, MBI, and sphericity. Variance analyses revealed a consistent and significant reduction in nearly all metrics in all residual tumors relative to TN (Fig. S9 & S10). As a result of the extensive mITH, we displayed a representative subset of the top, middle, and bottom 10% of mitochondria based on volume within each tumor. To visualize the mitochondrial structural diversity, we performed mito-otyping^27^, a karyotype-like arrangement of mitochondria to visualize complexity relative to volume (Fig. 4a, Fig. S13a). Consistent with the quantitative data, mito-otyping revealed substantial mITH in all tumors, along with significant shifts in the size and complexity of mitochondria following chemotherapy. Overall, these findings suggest that mitochondria tend to become less branched and/or elongated after treatment with DTX+CRB or AC. In contrast, single-agent DTX or CRB treatment produced divergent phenotypes between the two PDX models. As with the other mitochondrial features, all treatments elicited a substantial reduction in mITH.

**Figure 4.**
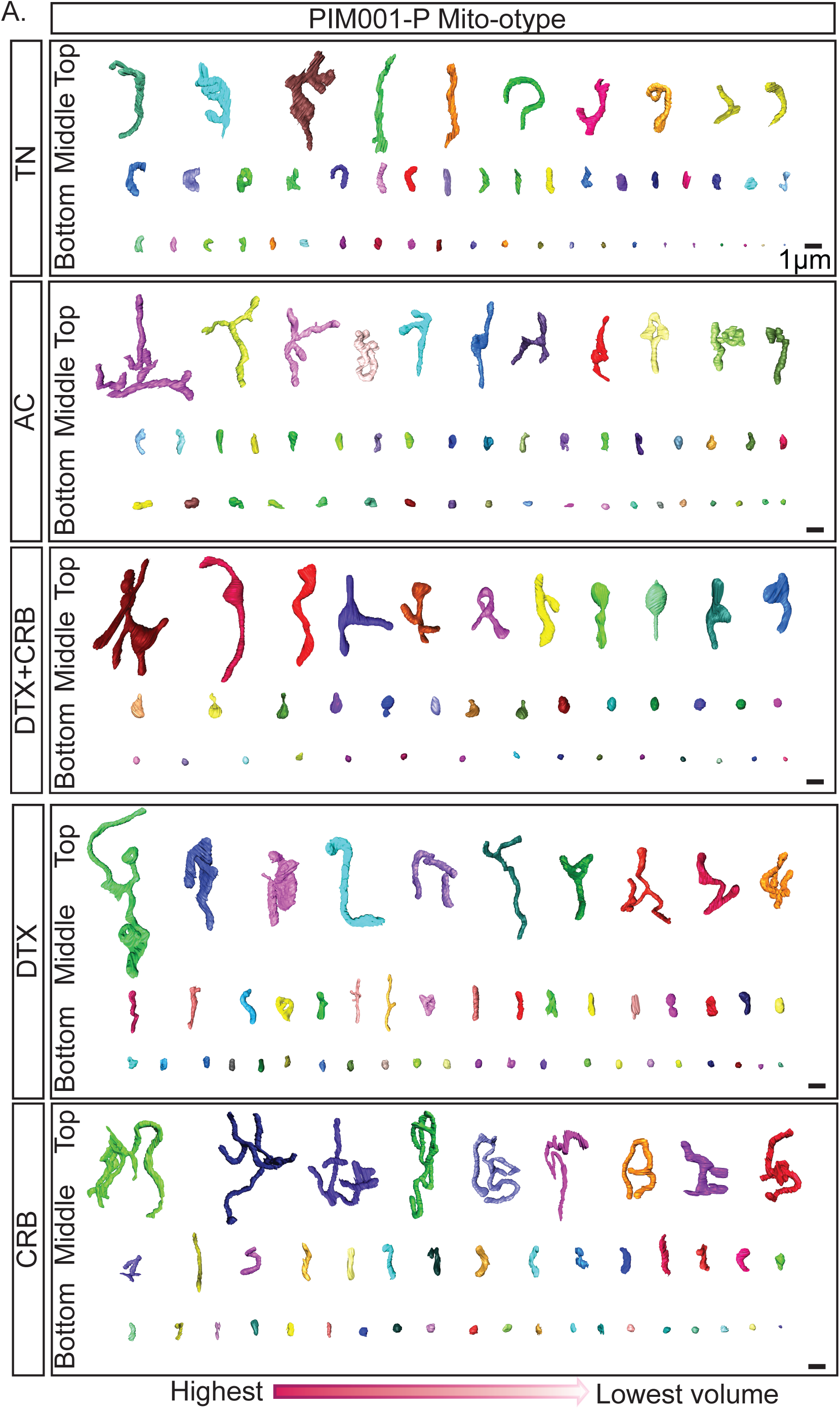
PIM001-P mito-otyping of the diverse mitochondrial structures within each treatment group. (a) Representative mitochondria structures from the bottom, middle, and top 10% of the 3D volume for tumors are displayed. Scale bars are 1µm.

### Mitochondrial-lipid droplet contacts increase in frequency while decrease in size upon chemotherapy

Lastly, while segmenting mitochondria in the second PDX model, WHIM14, we observed a striking abundance of lipid droplets (LDs). Since mitochondria do not function independently but instead interact with other organelles to facilitate the exchange of biomolecules, we extended our characterization to model 3D structures of mitochondrial-LD contact (MLC) sites. To this end, we first segmented 200 lipid droplets from each WHIM14 tumor (supplementary Table 3). We found that volume, area, and perimeter of lipid droplets increased in residual tumors persisting after single-agent chemotherapy treatments compared to the TN tumor (Fig. S14). Of the 500 segmented mitochondria and 200 segmented LDs in each WHIM14 tumor, we found 105 MLCs in the TN tumor, 188 in the DTX tumor, 137 in the CRB tumor, and 167 in the DTX+CRB tumor. Representative 3D MLCs are shown from the side, top, and zoomed top views (Fig. 5a-d’’). When considering the proportion of the mitochondria in contact with LDs, we observed that all chemotherapy-treated groups have increased MLCs compared to the TN (Fig. 5e). We then assessed the size of these contacts by evaluating their surface area and volume from the two ROIs selected (Fig. 5f & g). For both ROIs, the MLC surface areas and volumes, per the total sum of the mitochondrial surface areas or volumes, were substantially lower in residual tumors compared to the TN tumor (Fig. 5h & i). Taken together, these analyses demonstrate that although chemotherapy increased the size of LDs and the number of mitochondria–lipid droplet contacts in residual tumors compared to TN, the surface area and volume of the MLCs was markedly reduced. These results suggest that chemotherapy substantially altered crosstalk between LDs and mitochondria, simultaneously increasing the likelihood of contact formation while limiting the physical size of the interactions.

**Figure 5.**
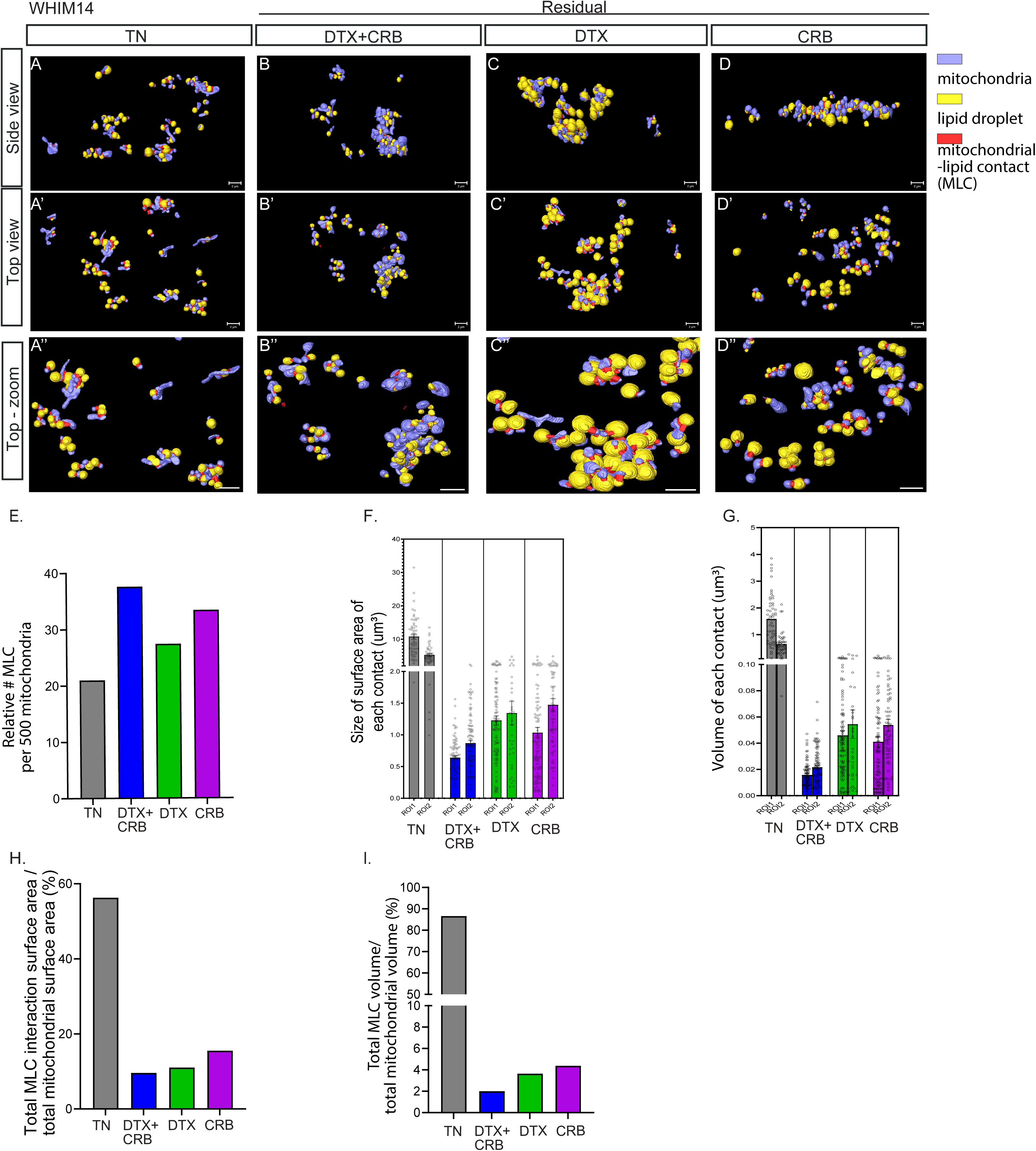
The frequency of MLCs increases upon chemotherapy, while the sizes of the contacts decrease in WHIM14. Quantification of how chemotherapy remodels mitochondrial-lipid droplet contacts (MLCs) for each treatment group: treatment-naïve (TN), Adriamycin + cyclophosphamide (AC), docetaxel + carboplatin (DTX+CRB), docetaxel (DTX), and carboplatin (CRB). (a-d’’) 3D rendering of the mitochondria in purple, lipid droplets in yellow, and their interaction in red. The scale bar is 2µm. The 3D renders show that TN exhibits relatively few but broad, sheet-like interfaces, whereas all treatment groups display numerous, smaller, punctate contacts scattered across organelles. Consistent with this visual shift, bar plots demonstrate that the fraction of MLCs among the total number of segmented mitochondria (E; % MLC per 500 mitochondria) is higher after treatment than in the TN group. In contrast, per-contact surface area (F) and volume (G) are largest in TN and significantly reduced in all treated conditions, indicating that individual interfaces between the mitochondria and LDs shrink following chemotherapy. This is also displayed in the bar plots, showing the total sum of either surface area (H) or volume of the MLC versus the total sum of each metric for the segmented mitochondria.

It is important to note that reduced surface area or volume of MLCs has been shown to reflect increased exchange of biomolecules such as ATP and fatty acids between mitochondria and LDs, potentially through closer physical connection^58,59,58^. In alignment with this, we previously published transcriptomic rewiring of the fatty acid metabolism pathway in residual PDX tumors persisting after AC treatment^13^. Further, proteogenomic analyses of 16 pairs of matched human TNBC biopsies prior to and following exposure to DTX+CRB revealed significant elevation of fatty acid metabolism pathways^41^. Together, these findings underscore the importance of future functional investigations into the role of mitochondria-lipid crosstalk in TNBC.

## DISCUSSION

Although chemotherapy is the backbone of TNBC treatment, there is no clear molecular rationale for choosing specific chemotherapeutic regimens and little understanding of the dependencies of chemo-refractory TNBC cells. While efforts have been made to identify molecular features that inform targeted treatments-such as with PARP inhibitors^2^ or immune checkpoint inhibitors^60^, poor understanding of how to select the most effective chemotherapies based on tumor molecular features remains. Thus, elucidating the biological properties of tumor cells that survive NACT is critically important for overcoming chemoresistance and improving overall survival for those 45% of TNBC patients whose tumors persist despite months of intensive, harsh chemotherapy treatments.

We previously established the functional importance of mitochondrial structure in supporting oxphos in chemoresistant TNBC. Furthermore, we demonstrated that inhibition of OXPHOS^13^ or of mitochondrial fusion^11^ in residual tumors persisting after chemotherapy significantly prolonged tumor control. Our study provides evidence that chemotherapy substantially alters the mitochondrial network in residual tumors across two unique PDX models treated with several conventional chemotherapeutic agents.

We observed greater volume, area, and perimeter of mitochondria in residual tumors after chemotherapy compared to treatment-naïve tumors in both TNBC PDX models. This may support increased metabolic capacity, as our previous study of TNBC demonstrated that mitochondrial elongation enhanced OXPHOS in residual tumors^11^. However, distinguishing the relative contributions of mitochondrial volume/area and the actual shape (elongated, spherical) remains a challenge in the mitochondrial structure field and is an active area of investigation. Thus, it is not immediately clear how the mitochondrial structures observed here may impact mitochondrial functions such as OXPHOS, redox regulation, anaplerosis, apoptosis, and cellular signaling. Mitochondrial dysfunction—an intricate condition with a wide ranging consequences—cannot be definitively inferred from the 3D mitochondrial structure alone. Nevertheless, given the importance of understanding mitochondrial dynamics in TNBC, the unique mitochondrial morphologies we observed in this study merit further functional investigations that may lead to therapeutic insights.

Mitochondria also interact and communicate with multiple organelles, such as the endoplasmic reticulum (ER). In fact, mitochondrial fission occurs at mitochondria-ER contact sites^21,61,62^. Mitochondria further engage with LDs, the nucleus, the Golgi, and lysosomes^22,63–66^. However, the prevalence and significance of these mitochondria—organelle contact sites remain hardly addressed in cancer and are practically unexplored in TNBC^67^. We observed an increase in mitochondrial surface area in residual tumors after chemotherapy, potentially creating more opportunities for organelle interactions. Indeed, one of our models exhibited marked levels of LDs, and we found that MLCs increased in all chemotherapy-treated tumors compared to the TN tumor. However, MLC size, measured by surface area and volume, was reduced in the residual tumors relative to TN. This suggests that while chemotherapy increases the frequency of MLCs, their smaller size may reflect tighter junctions between mitochondria and LDs^68^. LDs are evolutionarily conserved organelles known to support lipogenesis and triacyl glyceride synthesis^58,59^, while also storing and releasing lipids to meet the energetic demands of the cell in coordination with mitochondria^69,70^. Notably, peridroplet mitochondra, a subpopulation of the mitochondria tightly associated with LDs, are reported to increase under stress and display heightened activity. These mitochondria exhibit enhanced TCA cycling, electron transport chain complex activity, oxygen consumption rate, and ATP synthesis^53^. The physical interaction between mitochondria and LDs observed in this study may very well facilitate both mitochondrial activity and triacylglyceride synthesis, ultimately fueling the survival of residual cells. Further studies will be necessary to test this using an in vitro system that can be manipulated and imaged with proteins of interest tagged, which, unfortunately, is currently infeasible in PDX models. Nonetheless, mapping the connections between mitochondria and key organelles promises to provide critical insights into essential subcellular mechanisms of communication^7,21,69,71–74^.

We found that mitochondrial structural adaptations were more pronounced in tumors treated with a single-agent compared to those with a combination chemotherapies in PIM001-P, but not in WHIM14. Although combining chemotherapies or paring chemotherapies with other drugs can improve treatment efficacy, there is also compelling evidence of drug antagonism or weak additive effects^75–77^. In fact, our recent TNBC PDX study demonstrated that CRB or DTX alone were almost always more efficacious than their combination was, which only provided benefit in 9% of the models tested. Surprisingly, and quite concerningly, 12% of models tested exhibited antagonism between the two chemotherapies^40^. Therefore, it is crucial to continue in depth comparisons of conventional chemotherapy agents administered to TNBC patients every day. Our data indicate that the combining and scheduling of chemotherapies affects the ability of mitochondria to adapt, an insight that may be critical for the future success of mitochondria-targeting therapies developed for TNBC patients^78^.

We also observed extensive mITH, particularly in the TN tumors. Mitochondrial structural heterogeneity is tissue-specific and reflects the mitochondrial demands of the tissue. For example, skeletal muscle, which contains a mixture of mitochondria-rich and mitochondria-poor fibers, exhibits greater mitochondrial structural heterogeneity than cardiac muscle, which requires continuous ATP due to continuous contraction^27^. Similarly, mitochondria in neuronal axons and dendrites adopt different morphologies that match the differing energy demands of these neuronal compartments^30^. Thus, the mitochondrial ITH observed in TN tumors may parallel the spatial heterogeneity of proteins and other molecules reported in human TNBC specimens^57,79,80^. Heterogeneous mitochondria may provide energetic adaptability that supports cell survival under stressors, such as chemotherapy, although this remains to be experimentally tested. All chemotherapy treatments we examined markedly reduced mitochondrial ITH, suggesting that mitochondria may undergo a ‘bottleneck’ to sustain cell survival following chemotherapeutic assault. Previous studies in breast cancer have shown enrichment of a cancer stem cell-like, or ‘tumor-initiating cell,’ subpopulation after a variety of therapeutic exposures^81,82^. Another potential mechanism underpinning these changes is the dynamic adaptation of mitochondrial structure in response to chemotherapy, which may arise broadly across cells rather than being restricted to a subpopulation or clone. Substantial evidence supports this mechanism^8,13,83,84^ Indeed, mitochondria are highly dynamic organelles, with fusion and fission coordinated with multiple cellular processes including the cell cycle^7,21^.

Mitochondria are not static organelles; rather, cycles of fission and fusion support their dynamic nature^21,62,71^. Our imaging captured only a snapshot of mitochondrial structures in our tumor samples, which we suspect represents the ‘average’ mitochondrial state in each tumor specimen. As 3D methods with subcellular resolution advance to enable longitudinal or even real-time analyses, it may become possible to evaluate the rates of mitochondrial structural changes in experimental models^18,29,68,74,85–87^. As we showed previously, both pharmacologic and genetic perturbations can induce these mitochondrial structures^11^. It is reasonable to speculate that these structures are plastic and reversible, as reported prior in this PDX model, PIM001-P, treated with AC, where a reversible drug-tolerant state was observed^13^.

Our study has several limitations. While SBF-SEM provides detailed information on the 3D landscape of subcellular organelles^18^, it is exceptionally labor- and time-intensive, requiring specialized expertise^18^, which limits its routine use in research labs and in the clinic. For this reason, we performed an in-depth analysis of mitochondria for one tumor per treatment group, selecting spatially distinct ROIs for mitochondrial quantification to account for possible spatial heterogeneity within each tumor. It remains to be determined how reproducible mitochondrial features will be across large cohorts of human tumors. However, based on the similarities between the two single-agent groups as well as between the two combination groups, we suspect that our findings may be generalizable to a sizable subset of TNBC patients. Although efforts are underway to use AI or automate mitochondria and other organelle analyses, algorithms often need to be retrained on new data sets^88^. It is also critically important to expand these studies to additional TNBC specimens from other patients because inter-patient heterogeneity within TNBC is well documented at the DNA, RNA, and protein levels^48,80,89^. Increasing interest in including 3D modeling underscores its potential to provide a more comprehensive understanding of cancer progression and drug responses from morphological perspective^90,91^. Mitochondrial structural differences among TNBC subtypes have not yet been characterized but will be important in future research. Finally, PDX models lack a fully intact immune system and are highly sensitive to chemotherapy toxicities; thus, it will be essential to validate these findings in human tumor specimens with a fully intact microenvironment. Nonetheless, work from our group and many others have shown PDX models faithfully capture key features of human cancer, including ITH, metabolic phenotypes, and chemotherapeutic responses^33,92,93^.

## ACKNOWLEDGEMENTS

We thank the breast cancer patients who donated their biopsies for PDX models. The generation of the PDX model PIM001-P was supported by a generous gift from the Cazalot family and the MD Anderson Women’s Cancer Moonshot Program. Dr. Helen Piwnica-Worms provided the PIM001-P PDX model via a materials transfer agreement. The BCM Patient Derived Xenograft Core guided the treatment regimens for DTX, CRB, DTX+CRB funded by CPRIT Core Facility Award (RP220646) and P30 Cancer Center Support Grant (NCI-CA125123). Dr. Junegoo Lee and Ms. Emily Goff aided with animal experiments. Dr. Tao Wang provided guidance for statistical analyses. Ms. Janice Cowden provided patient research advocacy support. The Mayo Clinic Microscopy and Cell Analysis Core provided experimental technical support. H&E, Ku80, and human mitochondria staining of PDX tumor sections were conducted at the BCM Pathology Core and Lab, supported by the Breast Center and a variety of research grants awarded to its faculty, including one of nine Specialized Programs of Research Excellence (SPORE) in Breast Cancer granted by the National Institute of Health. S10 grant (1S10OD028671-01) for digital imaging of the IHC slides. Dr. George Miles aided with the histological review of IHC and H&E-stained slides.

## DATA AND CODE AVAILABILITY

Mitochondria images and raw data are available from the corresponding authors upon request. All computational codes used for statistical analyses are available https://github.com/audra-lane/berner-et-al-2024. The PDX model PIM001-P can be made available through a materials transfer agreement with The University of Texas MD Anderson Cancer Center. The PDX model WHIM14 can be made available through a materials transfer agreement with Dr. Shunquan Li

## AUTHOR CONTRIBUTIONS

MJB, HKB, AOH, and GVE were responsible for the overall study design, experimentation, data interpretation, and writing of the manuscript.

AL conducted statistical analyses under the supervision of GVE.

SWW aided in PDX model treatment studies under the supervision of GVE.

BR, ZV, BS, FZ, MK, LV, AGM, AP, AK, CV, PV, PK, AC, DP, JSR, and ES aided in mitochondrial segmentation, analysis, and 3D rendering of figures and videos under the supervision of AOH.

MLB aided in PDX model treatment study design, implementation, and mitochondrial imaging under the supervision of GVE.

LED aided in the chemotherapeutic regimen design and dosing of PDX mice under the supervision of MTL.

## Supplemental Materials

**Supplementary Table 1. PIM001-P selected segmented mitochondria used for analyses**

The number of segmented mitochondria characterized during this study and the process for selecting 350 mitochondria per treatment for analysis.

**Supplementary Table 2. WHIM14 selected segmented mitochondria used for analyses**

The number of segmented mitochondria characterized during this study of the second PDX model, WHIM14. Two ROIs from one residual tumor of each treatment arm were segmented for a minimum of 250 mitochondria each. Random number generator was used to select 500 segmented for each treatment arm to be used in the analysis.

**Supplementary Table 3. WHIM14 selected segmented lipid droplets used for analyses**

Number of segmented lipid droplets characterized during this study of the second PDX model, WHIM14 for the lipid-mitochondrial interaction analyses. Two ROIs from one residual tumor of each treatment arm were segmented for a minimum of 100 mitochondria each. Random number generator was used to select 200 segmented for each treatment arm to be used in the analysis.

**Supplementary Figure 1. PDX TNBC tumors persist following conventional chemotherapy treatments in WHIM14**

(a) Tumor volumes were monitored biweekly following administration of chemotherapies to mice bearing orthotopic WHIM14 tumors (n = 6 mice/group). Arrows at the top indicate when chemotherapy was administered. Asterisks (*) above the x-axis indicate early euthanasia due to animal health concerns. Residual tumors were harvested, as noted by arrows. Error bars represent the standard error of the mean. (B) Residual tumors were harvested, processed for FFPE, and analyzed by H&E staining, IHC for Ku80, and human-specific mitochondria. Scale bars are 50 µm.

**Supplementary Figure 2. PIM001-P PDX tumors treated with one cycle of DTX+CRB avoid toxicity and persist**

(a) Normalized tumor volume from the day of starting treatment of PIM001-P treated with one dose of DTX+CRB and monitored biweekly (n = 2 TN; n=4 DTX+CRB). Treatment groups were treatment-naïve (TN) and docetaxel + carboplatin (DTX+CRB).

**Supplementary Figure 3. Several mitochondrial features are significantly increased after single-agent chemotherapy treatments in WHIM14**

(a-d) Representative SBF-SEM orthoslices. (a’-d’) Representative 3D SBF-SEM orthoslices with overlays of mitochondria segmentations. (a’’-d’’) Representative 3D SBF-SEM mitochondria segmentations for each treatment group: treatment-naïve (TN), docetaxel + carboplatin (DTX+CRB), docetaxel (DTX), and carboplatin (CRB). The scale bar is 1 µm. Mitochondrial measurements were calculated using Amira software for 500 segmented mitochondria (represented as dots) from each tumor for (e) volume, (f) 3D area, and (g) perimeter. Adjusted p-values (*p < 0.05, **p < 0.01, ***p < 0.001, ****p<0.0001) are shown on box and whisker plots as determined using the Mann–Whitney U test followed by the Holm method.

**Supplementary Figure 4. PIM001-P mitochondrial measurements across treatment groups do not have normal distributions**

(a) Q–q plots of the raw measurements across all tumors are shown. Each blue dot represents one of the segmented mitochondria from a total of 350 segmented mitochondria. The red line indicates a reference for where the normal distribution falls on the graph.

**Supplementary Figure 5. PIM001-P distribution of mitochondrial metrics within each treatment group**

Density plots are of log10 transformed data showing the distribution of the respective metric for the segmented mitochondria. The line indicates the median for treatment-naïve tumors. Treatment groups: treatment-naïve (TN), Adriamycin + cyclophosphamide (AC), docetaxel + carboplatin (DTX+CRB), docetaxel (DTX), and carboplatin (CRB).

**Supplementary Figure 6. PIM001-P single chemotherapeutic agent treated mitochondria are greater in width and length**

Mitochondrial measurements for width and length were calculated using Amira software for 350 segmented mitochondria (represented as dots) from each treatment group. Treatment groups: treatment-naïve (TN), Adriamycin + cyclophosphamide (AC), docetaxel + carboplatin (DTX+CRB), docetaxel (DTX), and carboplatin (CRB). (a) Width (b) Length. Adjusted p-values (*p < 0.05, **p < 0.01, ***p < 0.001, ****p<0.0001) are shown on box and whisker plots as determined using the Mann–Whitney U test followed by the Holm method. Density plots are of log10 transformed data showing the distribution of the respective metric for the segmented mitochondria. The line indicates the median for treatment-naïve tumors. (c) Density plot for width (d) Density plot for length.

**Supplementary Figure 7. WHIM14 distribution of mitochondrial metrics within each treatment group**

Density plots are of log10 transformed data showing the distribution of the respective metric for the segmented mitochondria. The line indicates the median for treatment-naïve tumors. Treatment groups: treatment-naïve (TN), docetaxel + carboplatin (DTX+CRB), docetaxel (DTX), and carboplatin (CRB).

**Supplementary Figure 8. Chemotherapy treatment increased the width and length of mitochondria in WHIM14**

Mitochondrial measurements for width and length were calculated using Amira software for 350 segmented mitochondria (represented as dots) from each treatment group. Treatment groups: treatment-naïve (TN), docetaxel + carboplatin (DTX+CRB), docetaxel (DTX), and carboplatin (CRB). (a) Width (b) Length. Adjusted p-values (*p < 0.05, **p < 0.01, ***p < 0.001, ****p<0.0001) are shown on box and whisker plots as determined using the Mann–Whitney U test followed by the Holm method. Density plots are of log10 transformed data showing the distribution of the respective metric for the segmented mitochondria. The line indicates the median for treatment-naïve tumors. (c) Density plot for width (d) Density plot for length. Treatment groups: treatment-naïve (TN), docetaxel + carboplatin (DTX+CRB), docetaxel (DTX), and carboplatin (CRB).

**Supplementary Figure 9. mITH was significantly reduced upon chemotherapy for several mitochondrial metrics in PIM001-P**

(a) Schematic for comparing the variance between treatment groups across all mitochondrial features. Treatment groups: treatment-naïve (TN), Adriamycin + cyclophosphamide (AC), docetaxel + carboplatin (DTX+CRB), docetaxel (DTX), and carboplatin (CRB). (b-i) Variance bar graphs for all 350 segmented mitochondrial volume, area, perimeter, width, length, mitochondrial branching index (MBI), sphericity, and mitochondrial complex index (MCI). F-test and Holm method for adjusted p-values (*p < 0.05, **p < 0.01, ***p < 0.001, ****p<0.0001).

**Supplementary Figure 10. WHIM14 treated with DTX+CRB or CRB only had a significant reduction in mITH**

Treatment groups: treatment-naïve (TN), docetaxel + carboplatin (DTX+CRB), docetaxel (DTX), and carboplatin (CRB). (a-h) Variance bar graphs for all 500 segmented mitochondrial volume, area, perimeter, width, length, mitochondrial branching index (MBI), sphericity, and mitochondrial complex index (MCI). F-test and Holm method for adjusted p-values (*p < 0.05, **p < 0.01, ***p < 0.001, ****p<0.0001).

**Supplementary Figure 11. WHIM14 residual tumors’ mitochondria are less branched and more spherical compared to TN tumors**

Mitochondrial networks revealed through 3D mitochondria rendered using SBF-SEM. (a) Representative longitudinal and transverse views of the segmented mitochondria for each treatment group: treatment-naïve (TN), docetaxel + carboplatin (DTX+CRB), docetaxel (DTX), and carboplatin (CRB). The scale bar is 1 µm. Mitochondrial measurements were calculated using Amira software for 500 segmented mitochondria (represented as dots) from each tumor for (b) sphericity (c) mitochondrial complex index (MCI), along with their respective method of measurements below. Adjusted p-values (*p < 0.05, **p < 0.01, ***p < 0.001, ****p<0.0001) are shown on box and whisker plots as determined using the Mann–Whitney U test followed by the Holm method.

**Supplementary Figure 12. MBI of PIM001-P and WHIM14**

Mitochondrial branching index (MBI). (a) diagram of MBI calculation. MBI of the segmented mitochondria for each treatment group: treatment-naïve (TN), Adriamycin + cyclophosphamide (AC), docetaxel + carboplatin (DTX+CRB), docetaxel (DTX), and carboplatin (CRB) for the two PDX models in the study. (b) PIM001-PI (c) WHIM14 Adjusted p-values (*p < 0.05, **p < 0.01, ***p < 0.001, ****p<0.0001) are shown on box and whisker plots as determined using the Mann–Whitney U test followed by the Holm method.

**Supplementary Figure 13. WHIM14 mito-otyping of the diverse mitochondrial structures observed within each treatment group**

(a) Representative mitochondria structures from the bottom, middle, and top 10% of the 3D volume for tumors are displayed. Scale bars are 0.2µm.

**Supplementary Figure 14. WHIM14 lipid droplet sizes increase after single-agent chemotherapy treatment**

Lipid droplet segmentation of WHIM14 PDX models from each treatment group: treatment-naïve (TN), docetaxel + carboplatin (DTX+CRB), docetaxel (DTX), and carboplatin (CRB). Lipid droplet measurements were calculated using Amira software for 200 segmented lipid droplets (represented as dots) from each tumor for (a) volume, (b) 3D area, and (c) perimeter (d) sphericity (e) lipid droplet complexity index. Adjusted p-values (*p < 0.05, **p < 0.01, ***p < 0.001, ****p<0.0001) are shown on box and whisker plots as determined using the Mann–Whitney U test followed by the Holm method.

## REFERENCES CITED

1 Harbeck, N. et al. Breast cancer. Nat Rev Dis Primers 5, 66 (2019). 10.1038/s41572-019-0111-2

2 Bianchini, G., De Angelis, C., Licata, L. & Gianni, L. Treatment landscape of triple-negative breast cancer - expanded options, evolving needs. Nat Rev Clin Oncol 19, 91–113 (2022). 10.1038/s41571-021-00565-2

3 Schmid, P. et al. Pembrolizumab for Early Triple-Negative Breast Cancer. N Engl J Med 382, 810–821 (2020). 10.1056/NEJMoa1910549

4 Vagia, E., Mahalingam, D. & Cristofanilli, M. The Landscape of Targeted Therapies in TNBC. Cancers (Basel) 12 (2020). 10.3390/cancers12040916

5 Symmans, W. F. et al. Long-Term Prognostic Risk After Neoadjuvant Chemotherapy Associated With Residual Cancer Burden and Breast Cancer Subtype. J Clin Oncol 35, 1049–1060 (2017). 10.1200/JCO.2015.63.1010

6 Yau, C. et al. Residual cancer burden after neoadjuvant chemotherapy and long-term survival outcomes in breast cancer: a multicentre pooled analysis of 5161 patients. Lancet Oncol 23, 149–160 (2022). 10.1016/S1470-2045(21)00589-1

7 Jenkins, B. C. et al. Mitochondria in disease: changes in shapes and dynamics. Trends Biochem Sci 49, 346–360 (2024). 10.1016/j.tibs.2024.01.011

8 Pendleton, K. E., Wang, K. & Echeverria, G. V. Rewiring of mitochondrial metabolism in therapy-resistant cancers: permanent and plastic adaptations. Front Cell Dev Biol 11, 1254313 (2023). 10.3389/fcell.2023.1254313

9 Zhao, J. et al. Mitochondrial dynamics regulates migration and invasion of breast cancer cells. Oncogene 32, 4814–4824 (2013). 10.1038/onc.2012.494

10 Humphries, B. A. et al. Enhanced mitochondrial fission suppresses signaling and metastasis in triple-negative breast cancer. Breast Cancer Res 22, 60 (2020). 10.1186/s13058-020-01301-x

11 Baek, M. L. et al. Mitochondrial structure and function adaptation in residual triple negative breast cancer cells surviving chemotherapy treatment. Oncogene 42, 1117–1131 (2023). 10.1038/s41388-023-02596-8

12 Herkenne, S. et al. Developmental and Tumor Angiogenesis Requires the Mitochondria-Shaping Protein Opa1. Cell Metab 31, 987–1003 e1008 (2020). 10.1016/j.cmet.2020.04.007

13 Echeverria, G. V. et al. Resistance to neoadjuvant chemotherapy in triple-negative breast cancer mediated by a reversible drug-tolerant state. Sci Transl Med 11 (2019). 10.1126/scitranslmed.aav0936

14 Herkenne, S. et al. Developmental and Tumor Angiogenesis Requires the Mitochondria-Shaping Protein Opa1. Cell Metab 31, 987–1003.e1008 (2020). 10.1016/j.cmet.2020.04.007

15 Pellattiero, A. et al. Small molecule OPA1 inhibitors amplify cytochrome c release and reverse cancer cells resistance to Bcl-2 inhibitors. Sci Adv 11, eadx4562 (2025). 10.1126/sciadv.adx4562

16 Denk, W. & Horstmann, H. Serial block-face scanning electron microscopy to reconstruct three-dimensional tissue nanostructure. PLoS Biol 2, e329 (2004). 10.1371/journal.pbio.0020329

17 Jiao, W., Chatton, J. Y. & Genoud, C. Mitochondria morphometry in 3D datasets obtained from mouse brains with serial block-face scanning electron microscopy. Methods Cell Biol 177, 197–211 (2023). 10.1016/bs.mcb.2023.01.021

18 Marshall, A. G. et al. Serial Block Face-Scanning Electron Microscopy as a Burgeoning Technology. Adv Biol (Weinh) 7, e2300139 (2023). 10.1002/adbi.202300139

19 Garza-Lopez, E. et al. Correction: Garza-Lopez, et al. Protocols for Generating Surfaces and Measuring 3D Organelle Morphology Using Amira. Cells 2022, 11, 65. Cells 12 (2023). 10.3390/cells12101356

20 Garza-Lopez, E. et al. Protocols for Generating Surfaces and Measuring 3D Organelle Morphology Using Amira. Cells 11 (2021). 10.3390/cells11010065

21 Glancy, B., Kim, Y., Katti, P. & Willingham, T. B. The Functional Impact of Mitochondrial Structure Across Subcellular Scales. Front Physiol 11, 541040 (2020). 10.3389/fphys.2020.541040

22 Beasley, H. K., Rodman, T. A., Collins, G. V., Hinton, A., Jr. & Exil, V. TMEM135 is a Novel Regulator of Mitochondrial Dynamics and Physiology with Implications for Human Health Conditions. Cells 10 (2021). 10.3390/cells10071750

23 Hinton, A., Jr., et al. A Comprehensive Approach to Sample Preparation for Electron Microscopy and the Assessment of Mitochondrial Morphology in Tissue and Cultured Cells. Adv Biol (Weinh) 7, e2200202 (2023). 10.1002/adbi.202200202

24 Neikirk, K. et al. Systematic Transmission Electron Microscopy-Based Identification and 3D Reconstruction of Cellular Degradation Machinery. Adv Biol (Weinh) 7, e2200221 (2023). 10.1002/adbi.202200221

25 Chen, L. et al. Positive feedback loop between mitochondrial fission and Notch signaling promotes survivin-mediated survival of TNBC cells. Cell Death Dis 9, 1050 (2018). 10.1038/s41419-018-1083-y

26 Si, L. et al. Silibinin inhibits migration and invasion of breast cancer MDA-MB-231 cells through induction of mitochondrial fusion. Mol Cell Biochem 463, 189–201 (2020). 10.1007/s11010-019-03640-6

27 Vincent, A. E. et al. Quantitative 3D Mapping of the Human Skeletal Muscle Mitochondrial Network. Cell Rep 26, 996–1009 e1004 (2019). 10.1016/j.celrep.2019.01.010

28 Vue, Z. et al. Three-dimensional mitochondria reconstructions of murine cardiac muscle changes in size across aging. Am J Physiol Heart Circ Physiol 325, H965–H982 (2023). 10.1152/ajpheart.00202.2023

29 Vue, Z. et al. 3D reconstruction of murine mitochondria reveals changes in structure during aging linked to the MICOS complex. Aging Cell 22, e14009 (2023). 10.1111/acel.14009

30 Faitg, J. et al. 3D neuronal mitochondrial morphology in axons, dendrites, and somata of the aging mouse hippocampus. Cell Rep 36, 109509 (2021). 10.1016/j.celrep.2021.109509

31 Jadav, N. et al. Beyond the surface: Investigation of tumorsphere morphology using volume electron microscopy. J Struct Biol 215, 108035 (2023). 10.1016/j.jsb.2023.108035

32 Han, M. et al. Spatial mapping of mitochondrial networks and bioenergetics in lung cancer. Nature 615, 712–719 (2023). 10.1038/s41586-023-05793-3

33 Bruna, A. et al. A Biobank of Breast Cancer Explants with Preserved Intra-tumor Heterogeneity to Screen Anticancer Compounds. Cell 167, 260–274 e222 (2016). 10.1016/j.cell.2016.08.041

34 Dobrolecki, L. E. et al. Patient-derived xenograft (PDX) models in basic and translational breast cancer research. Cancer Metastasis Rev 35, 547–573 (2016). 10.1007/s10555-016-9653-x

35 Echeverria, G. V. et al. High-resolution clonal mapping of multi-organ metastasis in triple negative breast cancer. Nat Commun 9, 5079 (2018). 10.1038/s41467-018-07406-4

36 Eirew, P. et al. Dynamics of genomic clones in breast cancer patient xenografts at single-cell resolution. Nature 518, 422–426 (2015). 10.1038/nature13952

37 Savage, P. et al. Chemogenomic profiling of breast cancer patient-derived xenografts reveals targetable vulnerabilities for difficult-to-treat tumors. Commun Biol 3, 310 (2020). 10.1038/s42003-020-1042-x

38 Baek, M., Chang, J. T. & Echeverria, G. V. Methodological Advancements for Investigating Intra-tumoral Heterogeneity in Breast Cancer at the Bench and Bedside. J Mammary Gland Biol Neoplasia 25, 289–304 (2020). 10.1007/s10911-020-09470-3

39 Eccles, S. A. et al. Critical research gaps and translational priorities for the successful prevention and treatment of breast cancer. Breast Cancer Res 15, R92 (2013). 10.1186/bcr3493

40 Lei, J. T. et al. Patient-Derived Xenografts of Triple-Negative Breast Cancer Enable Deconvolution and Prediction of Chemotherapy Responses. bioRxiv (2025). 10.1101/2024.12.09.627518

41 Anurag, M. et al. Proteogenomic Markers of Chemotherapy Resistance and Response in Triple-Negative Breast Cancer. Cancer Discov 12, 2586–2605 (2022). 10.1158/2159-8290.CD-22-0200

42 Virtanen, P. et al. SciPy 1.0: fundamental algorithms for scientific computing in Python. Nat Methods 17, 261–272 (2020). 10.1038/s41592-019-0686-2

43 Seabold, S., and Perktold, J. . Statsmodels: Econometric and statistical modeling with python. Proceedings of the 9th python in science conference (2010). 10.25080/Majora-92bf1922-011

44 McKinney, W. Data structures for statistical computing in python. (2010).

45 Team, R. C. R: A Language and Environment for Statistical Computing, <<https://www.R-project.org/>> (2023).

46 Li, S. et al. Endocrine-therapy-resistant ESR1 variants revealed by genomic characterization of breast-cancer-derived xenografts. Cell Rep 4, 1116–1130 (2013). 10.1016/j.celrep.2013.08.022

47 Fisher, E. R. et al. Pathobiology of preoperative chemotherapy: findings from the National Surgical Adjuvant Breast and Bowel (NSABP) protocol B-18. Cancer 95, 681–695 (2002). 10.1002/cncr.10741

48 Shah, S. P. et al. The clonal and mutational evolution spectrum of primary triple-negative breast cancers. Nature 486, 395–399 (2012). 10.1038/nature10933

49 Wang, Y. et al. Clonal evolution in breast cancer revealed by single nucleus genome sequencing. Nature 512, 155–160 (2014). 10.1038/nature13600

50 Nik-Zainal, S. et al. The life history of 21 breast cancers. Cell 149, 994–1007 (2012). 10.1016/j.cell.2012.04.023

51 Risom, T. et al. Differentiation-state plasticity is a targetable resistance mechanism in basal-like breast cancer. Nat Commun 9, 3815 (2018). 10.1038/s41467-018-05729-w

52 Kim, C. et al. Chemoresistance Evolution in Triple-Negative Breast Cancer Delineated by Single-Cell Sequencing. Cell 10.1016/j.cell.2018.03.041

53 Li, X. et al. Intrinsic resistance of tumorigenic breast cancer cells to chemotherapy. J Natl Cancer Inst 100, 672–679 (2008). 10.1093/jnci/djn123

54 Ali, H. R. et al. Imaging mass cytometry and multiplatform genomics define the phenogenomic landscape of breast cancer. Nature Cancer 1, 163–175 (2020). 10.1038/s43018-020-0026-6

55 Fischer, J. R. et al. Multiplex imaging of breast cancer lymph node metastases identifies prognostic single-cell populations independent of clinical classifiers. Cell Rep Med 4, 100977 (2023). 10.1016/j.xcrm.2023.100977

56 Gruosso, T. et al. Spatially distinct tumor immune microenvironments stratify triple-negative breast cancers. J Clin Invest 129, 1785–1800 (2019). 10.1172/JCI96313

57 Wagner, J. et al. A Single-Cell Atlas of the Tumor and Immune Ecosystem of Human Breast Cancer. Cell 177, 1330–1345 e1318 (2019). 10.1016/j.cell.2019.03.005

58 Benador, I. Y. et al. Mitochondria Bound to Lipid Droplets Have Unique Bioenergetics, Composition, and Dynamics that Support Lipid Droplet Expansion. Cell Metab 27, 869–885 e866 (2018). 10.1016/j.cmet.2018.03.003

59 Benador, I. Y., Veliova, M., Liesa, M. & Shirihai, O. S. Mitochondria Bound to Lipid Droplets: Where Mitochondrial Dynamics Regulate Lipid Storage and Utilization. Cell Metab 29, 827–835 (2019). 10.1016/j.cmet.2019.02.011

60 Cortes, J. et al. Pembrolizumab plus chemotherapy versus placebo plus chemotherapy for previously untreated locally recurrent inoperable or metastatic triple-negative breast cancer (KEYNOTE-355): a randomised, placebo-controlled, double-blind, phase 3 clinical trial. Lancet 396, 1817–1828 (2020). 10.1016/S0140-6736(20)32531-9

61 Hinton, A., Jr., et al. ATF4-dependent increase in mitochondrial-endoplasmic reticulum tethering following OPA1 deletion in skeletal muscle. J Cell Physiol 239, e31204 (2024). 10.1002/jcp.31204

62 Robertson, G. L. et al. DRP1 mutations associated with EMPF1 encephalopathy alter mitochondrial membrane potential and metabolic programs. J Cell Sci 136 (2023). 10.1242/jcs.260370

63 Filadi, R. et al. TOM70 Sustains Cell Bioenergetics by Promoting IP3R3-Mediated ER to Mitochondria Ca(2+) Transfer. Curr Biol 28, 369–382 e366 (2018). 10.1016/j.cub.2017.12.047

64 Giorgi, C., Marchi, S. & Pinton, P. The machineries, regulation and cellular functions of mitochondrial calcium. Nat Rev Mol Cell Biol 19, 713–730 (2018). 10.1038/s41580-018-0052-8

65 Szymanski, J. et al. Interaction of Mitochondria with the Endoplasmic Reticulum and Plasma Membrane in Calcium Homeostasis, Lipid Trafficking and Mitochondrial Structure. Int J Mol Sci 18 (2017). 10.3390/ijms18071576

66 Neikirk, K. et al. Call to action to properly utilize electron microscopy to measure organelles to monitor disease. Eur J Cell Biol 102, 151365 (2023). 10.1016/j.ejcb.2023.151365

67 Yu, H., Sun, C., Gong, Q. & Feng, D. Mitochondria-Associated Endoplasmic Reticulum Membranes in Breast Cancer. Front Cell Dev Biol 9, 629669 (2021). 10.3389/fcell.2021.629669

68 Crabtree, A. et al. Quantitative assessment of morphological changes in lipid droplets and lipid-mito interactions with aging in brown adipose. J Cell Physiol 239, e31340 (2024). 10.1002/jcp.31340

69 Jarc, E. & Petan, T. Lipid Droplets and the Management of Cellular Stress. Yale J Biol Med 92, 435–452 (2019).

70 Viale, A. et al. Oncogene ablation-resistant pancreatic cancer cells depend on mitochondrial function. Nature 514, 628–632 (2014). 10.1038/nature13611

71 Katti, P., Ajayi, P. T., Aponte, A., Bleck, C. K. E. & Glancy, B. Identification of evolutionarily conserved regulators of muscle mitochondrial network organization. Nat Commun 13, 6622 (2022). 10.1038/s41467-022-34445-9

72 Glancy, B. & Balaban, R. S. Role of mitochondrial Ca2+ in the regulation of cellular energetics. Biochemistry 51, 2959–2973 (2012). 10.1021/bi2018909

73 Lam, J. et al. A Universal Approach to Analyzing Transmission Electron Microscopy with ImageJ. Cells 10 (2021). 10.3390/cells10092177

74 Shao, B. et al. Ablation of Sam50 is associated with fragmentation and alterations in metabolism in murine and human myotubes. J Cell Physiol 239, e31293 (2024). 10.1002/jcp.31293

75 Osborne, C. K., Kitten, L. & Arteaga, C. L. Antagonism of chemotherapy-induced cytotoxicity for human breast cancer cells by antiestrogens. J Clin Oncol 7, 710–717 (1989). 10.1200/JCO.1989.7.6.710

76 Palmer, A. C., Chidley, C. & Sorger, P. K. A curative combination cancer therapy achieves high fractional cell killing through low cross-resistance and drug additivity. Elife 8 (2019). 10.7554/eLife.50036

77 Plana, D., Palmer, A. C. & Sorger, P. K. Independent Drug Action in Combination Therapy: Implications for Precision Oncology. Cancer Discov 12, 606–624 (2022). 10.1158/2159-8290.CD-21-0212

78 Bu, H. et al. Spinal IFN-gamma-induced protein-10 (CXCL10) mediates metastatic breast cancer-induced bone pain by activation of microglia in rat models. Breast Cancer Res Treat 143, 255–263 (2014). 10.1007/s10549-013-2807-4

79 Ali, H. R. et al. Imaging mass cytometry and multiplatform genomics define the phenogenomic landscape of breast cancer. Nat Cancer 1, 163–175 (2020). 10.1038/s43018-020-0026-6

80 Jackson, H. W. et al. The single-cell pathology landscape of breast cancer. Nature 578, 615–620 (2020). 10.1038/s41586-019-1876-x

81 Creighton, C. J. et al. Residual breast cancers after conventional therapy display mesenchymal as well as tumor-initiating features. Proc Natl Acad Sci U S A 106, 13820–13825 (2009). 10.1073/pnas.0905718106

82 Balko, J. M. et al. Profiling of residual breast cancers after neoadjuvant chemotherapy identifies DUSP4 deficiency as a mechanism of drug resistance. Nat Med 18, 1052–1059 (2012). 10.1038/nm.2795

83 Kim, C. et al. Chemoresistance Evolution in Triple-Negative Breast Cancer Delineated by Single-Cell Sequencing. Cell 173, 879–893 e813 (2018). 10.1016/j.cell.2018.03.041

84 Almendro, V. et al. Inference of tumor evolution during chemotherapy by computational modeling and in situ analysis of genetic and phenotypic cellular diversity. Cell Rep 6, 514–527 (2014). 10.1016/j.celrep.2013.12.041

85 Crabtree, A. et al. Defining Mitochondrial Cristae Morphology Changes Induced by Aging in Brown Adipose Tissue. Adv Biol (Weinh) 8, e2300186 (2024). 10.1002/adbi.202300186

86 Marshall, A. G., Krystofiak, E., Damo, S. M. & Hinton, A., Jr. Correlative light-electron microscopy: integrating dynamics to structure. Trends Biochem Sci 48, 826–827 (2023). 10.1016/j.tibs.2023.05.003

87 Scudese, E. et al. 3D Mitochondrial Structure in Aging Human Skeletal Muscle: Insights Into MFN-2-Mediated Changes. Aging Cell 24, e70054 (2025). 10.1111/acel.70054

88 Heinrich, L. et al. Whole-cell organelle segmentation in volume electron microscopy. Nature 599, 141–146 (2021). 10.1038/s41586-021-03977-3

89 Lehmann, B. D. et al. Identification of human triple-negative breast cancer subtypes and preclinical models for selection of targeted therapies. J Clin Invest 121, 2750–2767 (2011). 10.1172/JCI45014

90 Rozenblatt-Rosen, O. et al. The Human Tumor Atlas Network: Charting Tumor Transitions across Space and Time at Single-Cell Resolution. Cell 181, 236–249 (2020). 10.1016/j.cell.2020.03.053

91 Johnson, B. E. et al. An omic and multidimensional spatial atlas from serial biopsies of an evolving metastatic breast cancer. Cell Rep Med 3, 100525 (2022). 10.1016/j.xcrm.2022.100525

92 Petrosyan, V. et al. Identifying biomarkers of differential chemotherapy response in TNBC patient-derived xenografts with a CTD/WGCNA approach. iScience 26, 105799 (2023). 10.1016/j.isci.2022.105799

93 Guillen, K. P. et al. A human breast cancer-derived xenograft and organoid platform for drug discovery and precision oncology. Nat Cancer 3, 232–250 (2022). 10.1038/s43018-022-00337-6

