## Supplementary Tables 1-3 for "Three-dimensional electron microscopy uncovers mitochondrial network remodeling in residual triple negative breast cancer persisting after conventional chemotherapy treatments"

PDX Model: PIM001-P

| Treatment Naïve (TN) | ROI | #1 | #2 | #3 | #4 |
|---|---|---|---|---|---|
| | Number of mito annotated after QC | 142 | 254 | 53 | 53 |
| | Number of mito randomly selected for analysis from each ROI | 125 | 125 | 50 | 50 |
| Mouse #260 | TOTAL number of mito used for analysis | 350 | | | |

| Adriamycin and Cyclophosphamide (AC) | Set of mito segemented | #1 | #2 | #3 |
|---|---|---|---|---|
| | Number of mito annotated after QC | 262 | 59 | 50 |
| | Number of mito randomly selected for analysis from each ROI | 250 | 50 | 50 |
| Mouse #278 | TOTAL number of mito used for analysis | 350 | | |

| Docetaxel and Carboplatin (DTX+CRB) | Set of mito segemented | #1 | #2 | #3 |
|---|---|---|---|---|
| | Number of mito annotated after QC | 763 | 49 | 49 |
| | Number of mito randomly selected for analysis from each ROI | 252 | 49 | 49 |
| Mouse #270 | TOTAL number of mito used for analysis | 350 | | |

| Docetaxel (DTX) | Set of mito segemented | #1 | #2 | #3 |
|---|---|---|---|---|
| | Number of mito annotated after QC | 330 | 47 | 60 |
| | Number of mito randomly selected for analysis from each ROI | 253 | 47 | 50 |
| Mouse #257 | TOTAL number of mito used for analysis | 350 | | |

| Carboplatin (CRB) | Set of mito segemented | #1 | #2 | #3 |
|---|---|---|---|---|
| | Number of mito annotated after QC | 268 | 52 | 51 |
| | Number of mito randomly selected for analysis from each ROI | 250 | 50 | 50 |
| Mouse #268 | TOTAL number of mito used for analysis | 350 | | |

Supplementary Table 1. PIM001-P selected segmented mitochondria used for analyses.

PDX Model: WHIM14

| Treatment Naïve (TN) | ROI | #1 | #2 |
|---|---|---|---|
| | Number of mito annotated after QC | 250 | 250 |
| | Number of mito randomly selected for analysis from each ROI | 250 | 250 |
| Mouse #409 | analysis | 500 | |

| Docetaxel and Carboplatin (DTX+CRB) | ROI | #1 | #2 |
|---|---|---|---|
| | Number of mito annotated after QC | 252 | 250 |
| | Number of mito randomly selected for analysis from each ROI | 250 | 250 |
| Mouse #382 | analysis | 500 | |

| Docetaxel (DTX) | ROI | #1 | #2 |
|---|---|---|---|
| | Number of mito annotated after QC | 250 | 250 |
| | Number of mito randomly selected for analysis from each ROI | 250 | 250 |
| Mouse #390 | analysis | 500 | |

| Carboplatin (CRB) | ROI | #1 | #2 |
|---|---|---|---|
| | Number of mito annotated after QC | 255 | 255 |
| | Number of mito randomly selected for analysis from each ROI | 250 | 250 |
| Mouse #389 | [illegible] analysis | 500 | |

Supplementary Table 2. WHIM14 selected segmented mitochondria used for analyses.

PDX Model: WHIM14

| Treatment Naïve (TN) | ROI | #1 | #2 |
|---|---|---|---|
| | Number of mito annotated after QC | 169 | 114 |
| | Number of mito randomly selected for analysis from each ROI | 100 | 100 |
| Mouse #409 | analysis | 200 | |

| Docetaxel and Carboplatin (DTX+CRB) | ROI | #1 | #2 |
|---|---|---|---|
| | Number of mito annotated after QC | 109 | 150 |
| | Number of mito randomly selected for analysis from each ROI | 100 | 100 |
| Mouse #382 | analysis | 200 | |

| Docetaxel (DTX) | ROI | #1 | #2 |
|---|---|---|---|
| | Number of mito annotated after QC | 121 | 101 |
| | Number of mito randomly selected for analysis from each ROI | 100 | 100 |
| Mouse #390 | analysis | 200 | |

| Carboplatin (CRB) | ROI | #1 | #2 |
|---|---|---|---|
| | Number of mito annotated after QC | 118 | 103 |
| | Number of mito randomly selected for analysis from each ROI | 100 | 100 |
| Mouse #389 | analysis | 200 | |

Supplementary Table 3. WHIM14 selected segmented lipid droplets used for analyses.
