## Supplementary Figures 1-14 for "Three-dimensional electron microscopy uncovers mitochondrial network remodeling in residual triple negative breast cancer persisting after conventional chemotherapy treatments"

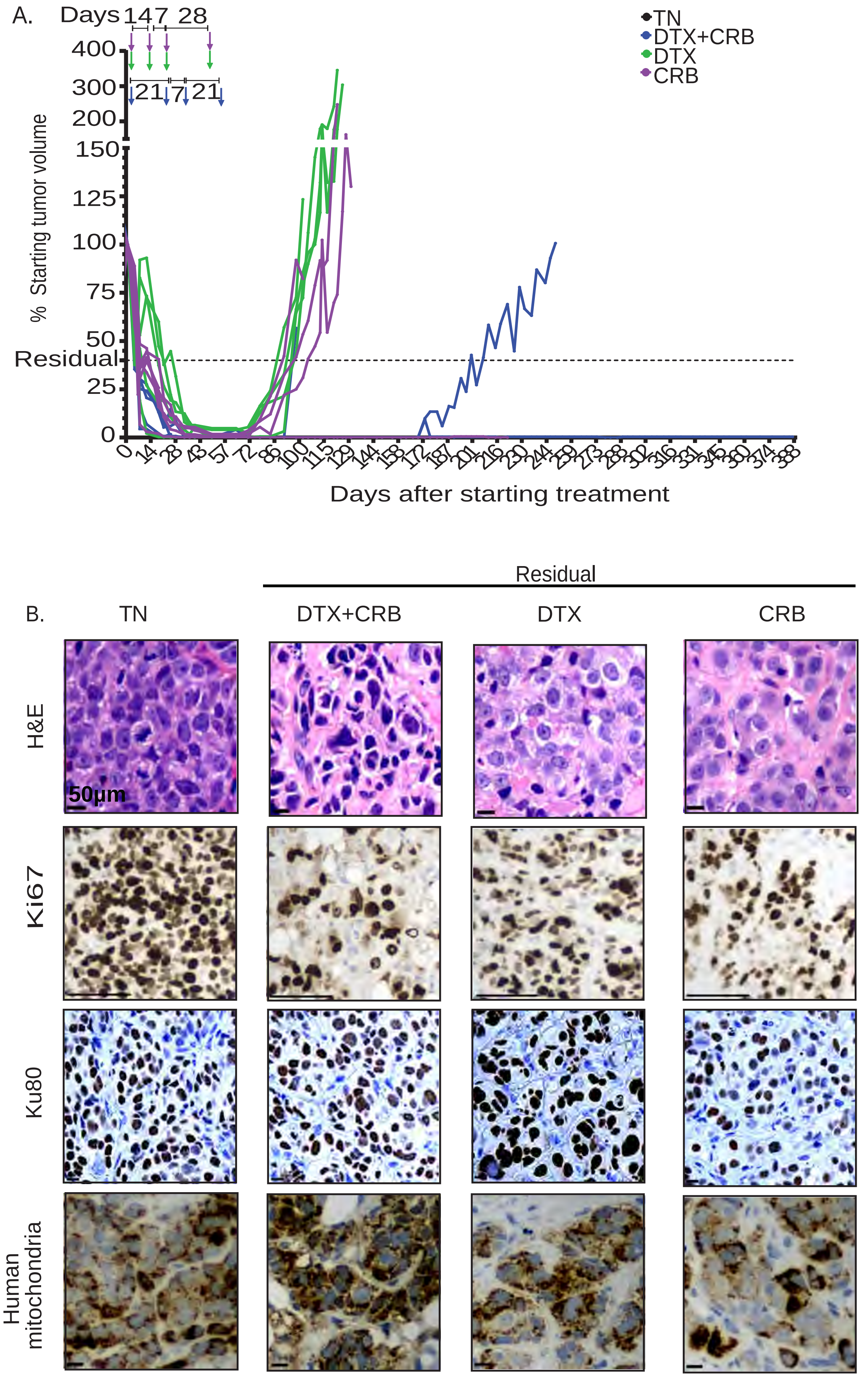


Figure S1. PDX TNBC tumors persist following conventional chemotherapy treatments in WHIM14.

A.

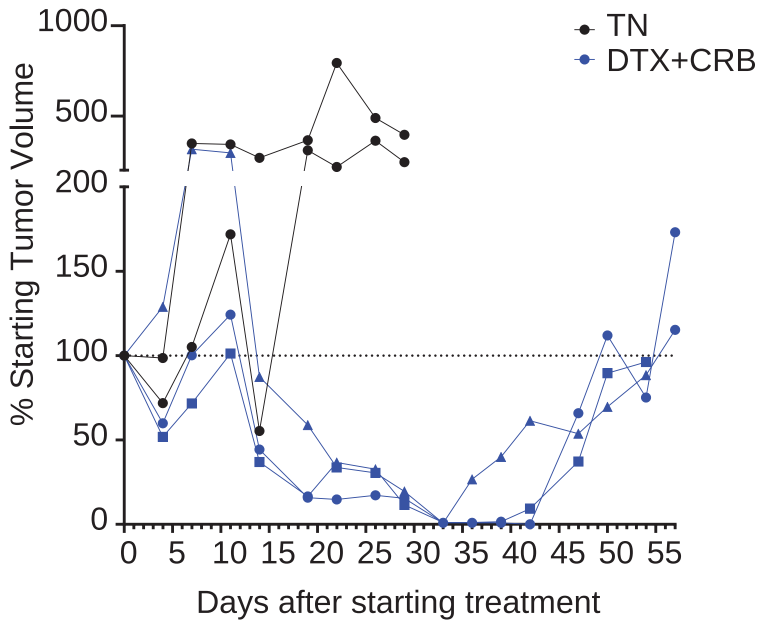


Figure S2. PIM001-P PDX tumors treated with one cycle of DTX+CRB avoid toxicity and persist.

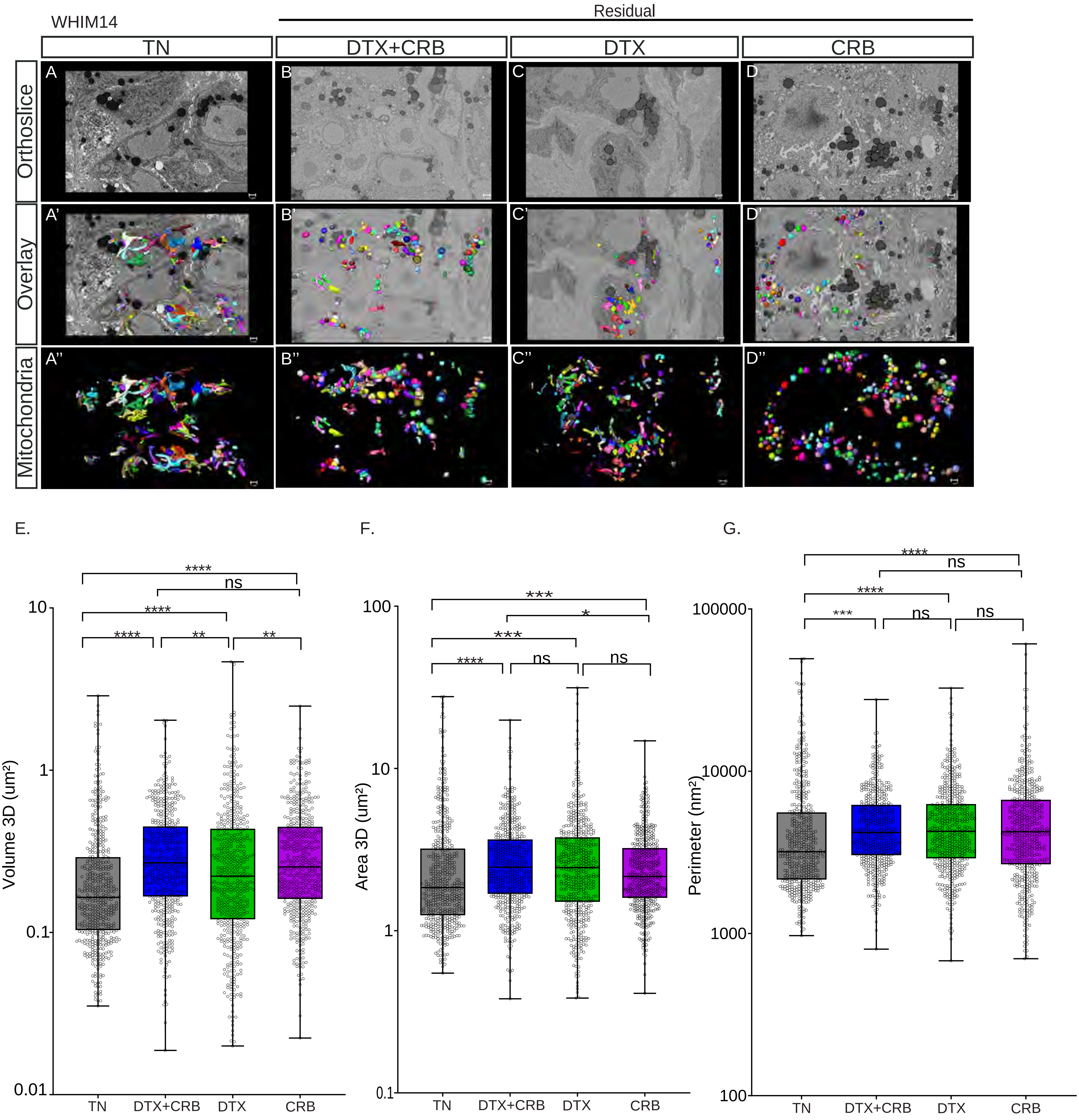


Figure S3. Several mitochondrial features are significantly increased after single-agent chemotherapy treatments in WHIM14.

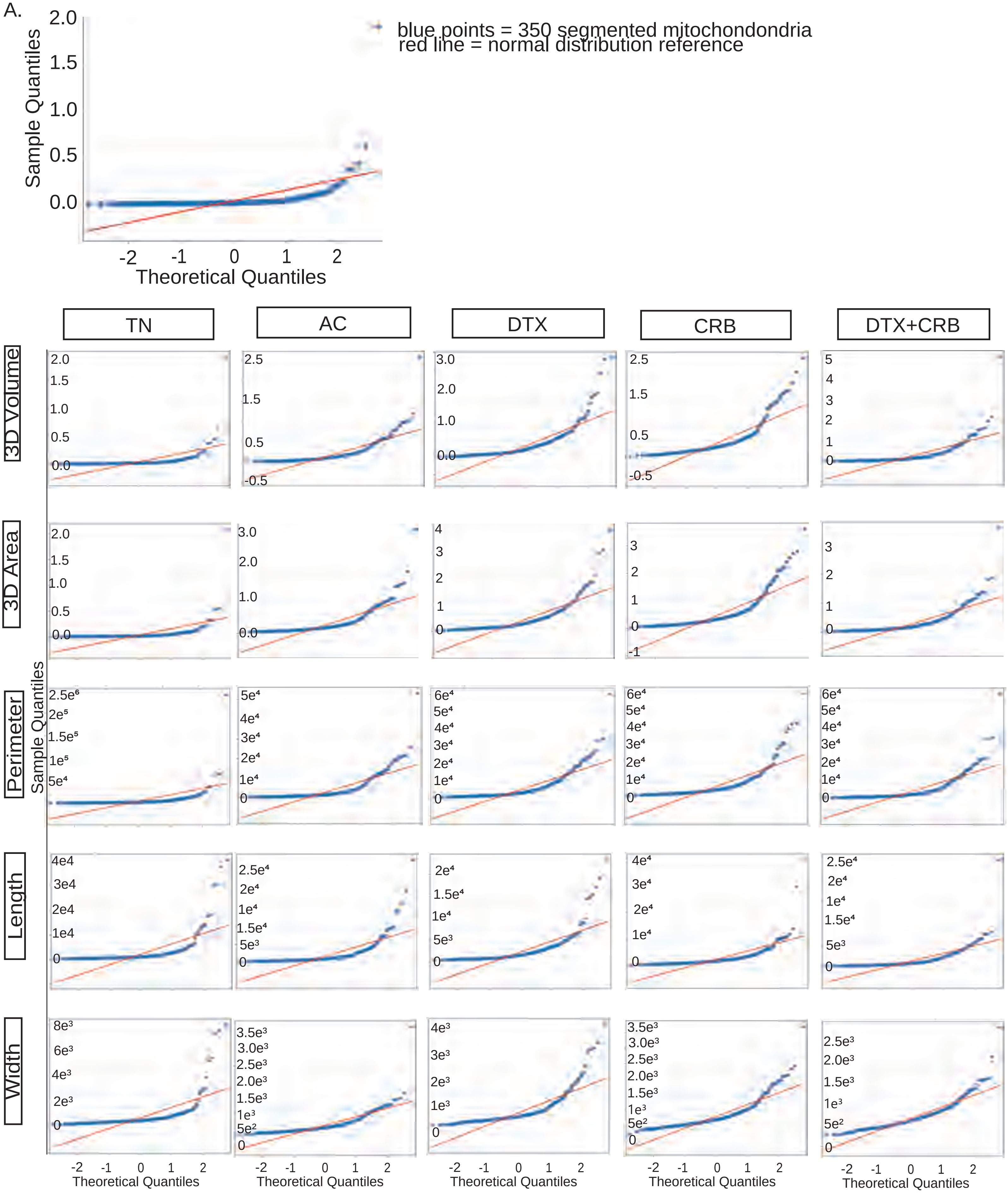


Figure S4. PIM001-P mitochondrial measurements across treatment groups do not have normal distributions.

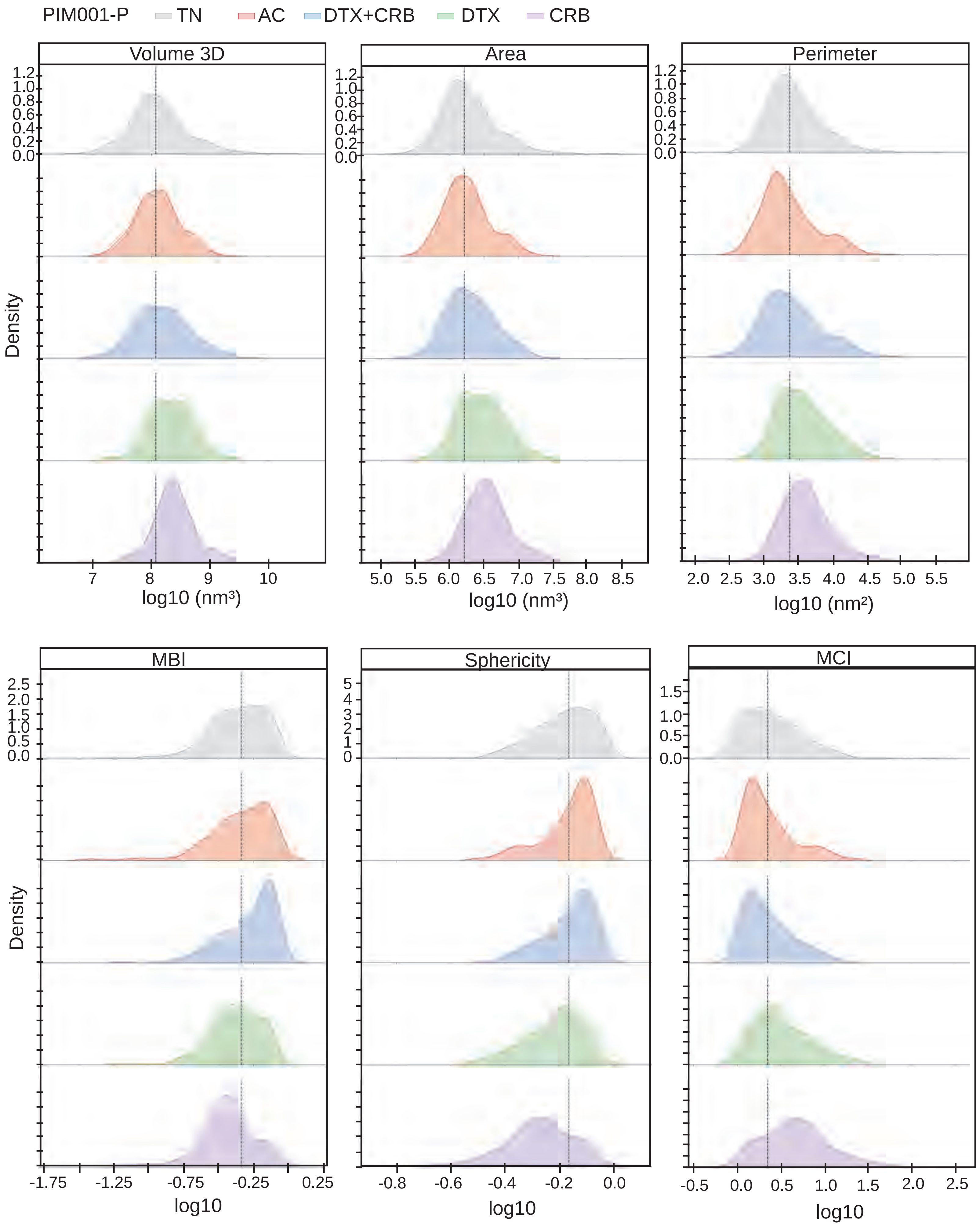


Figure S5. PIM001-P distribution of mitochondrial metrics within each treatment group.

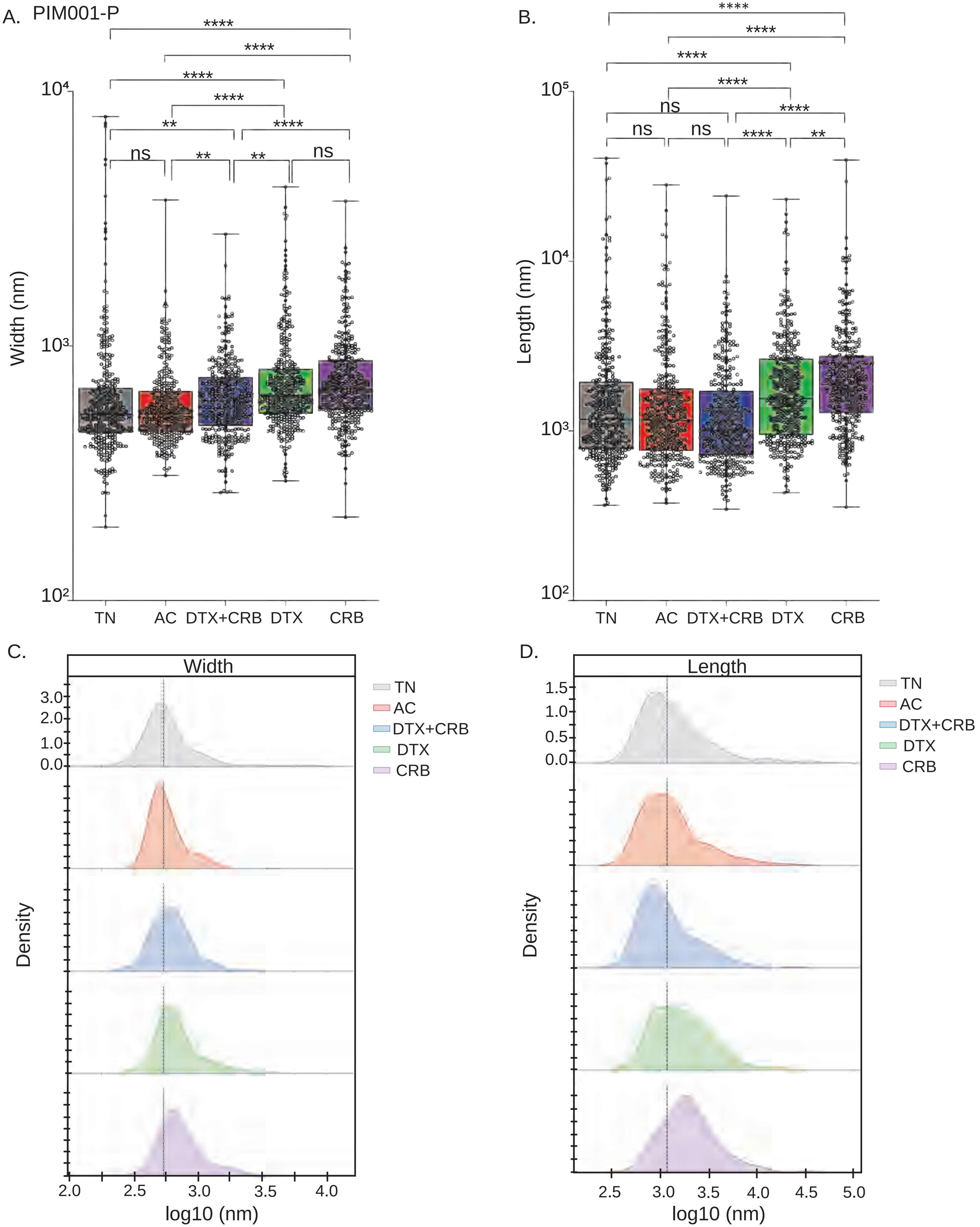


Figure S6. PIM001-P single chemotherapeutic agent treated mitochondria are greater in width and length.

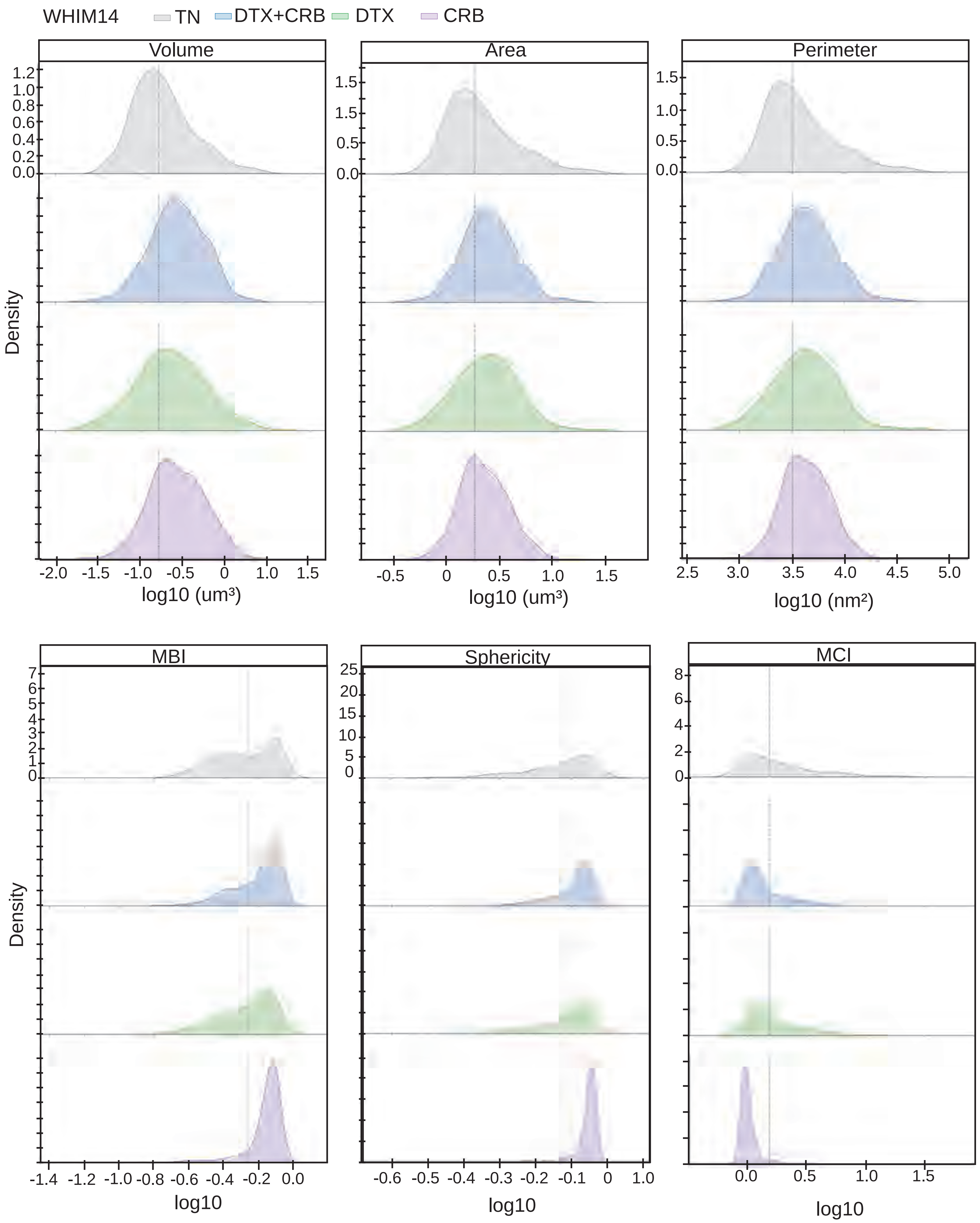


Figure S7. WHIM14 distribution of mitochondrial metrics within each treatment group.

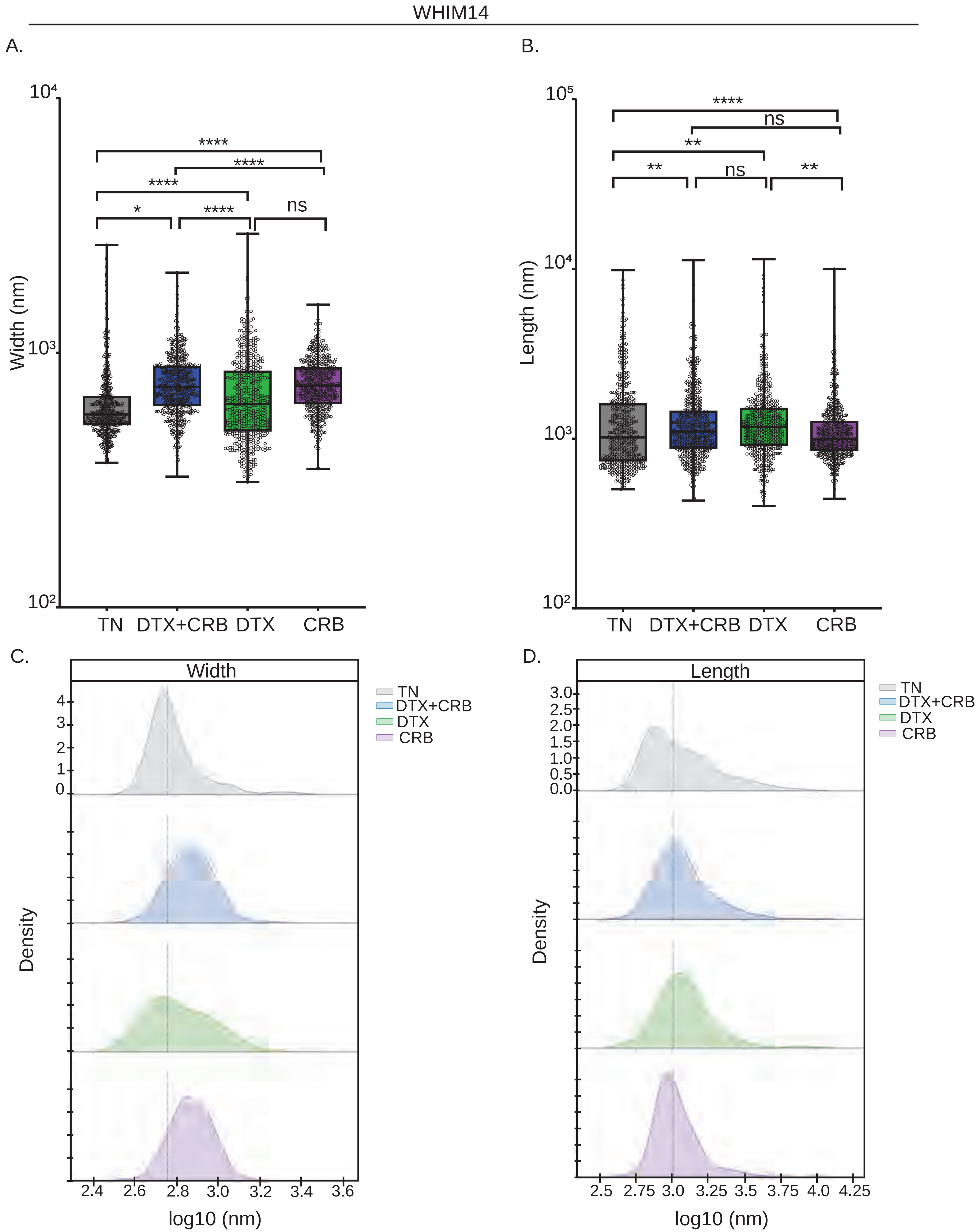


Figure S8. Chemotherapy treatment increases the width and length of mitochondria in the WHIM14.

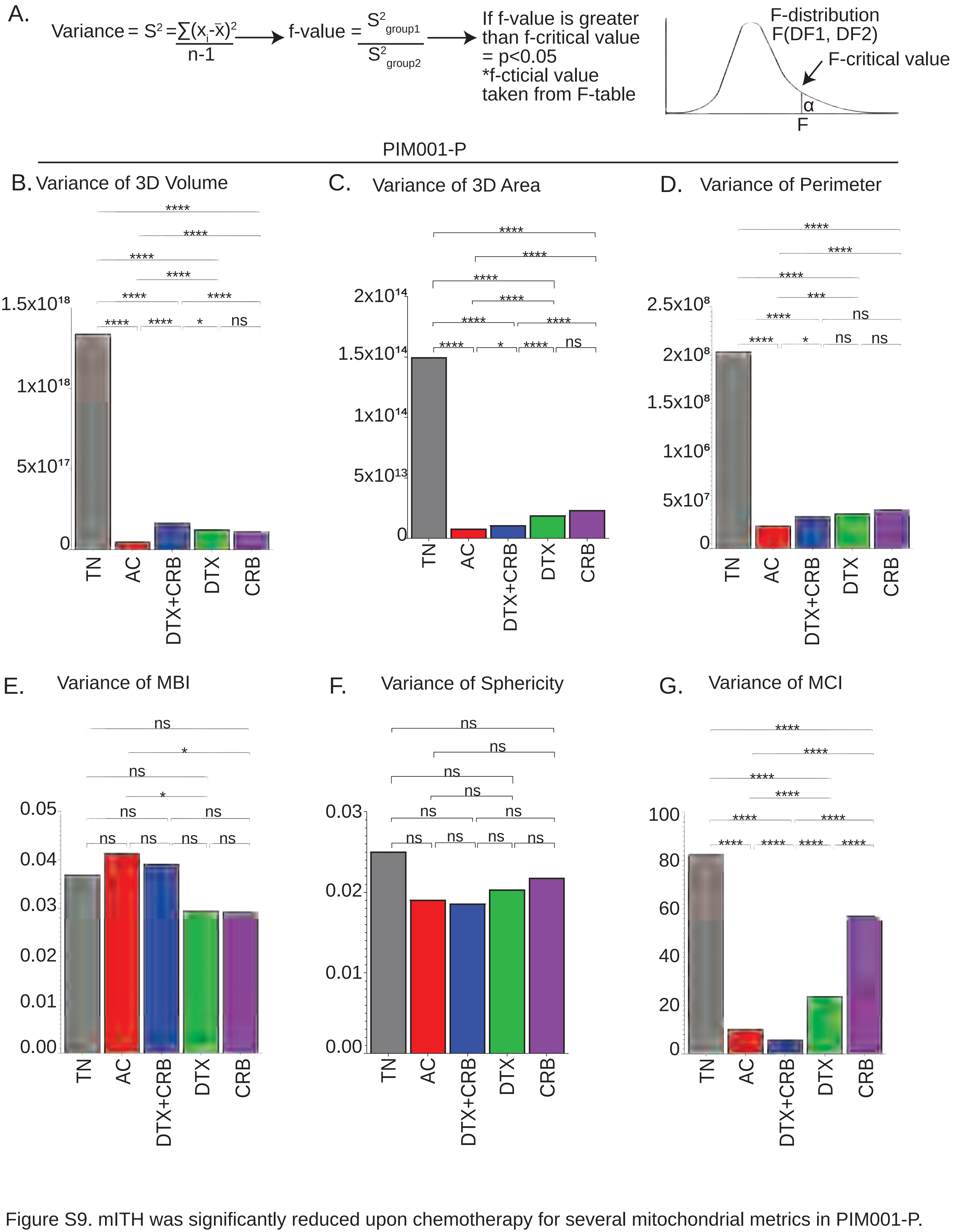


Figure S9. mITH was significantly reduced upon chemotherapy for several mitochondrial metrics in PIM001-P.

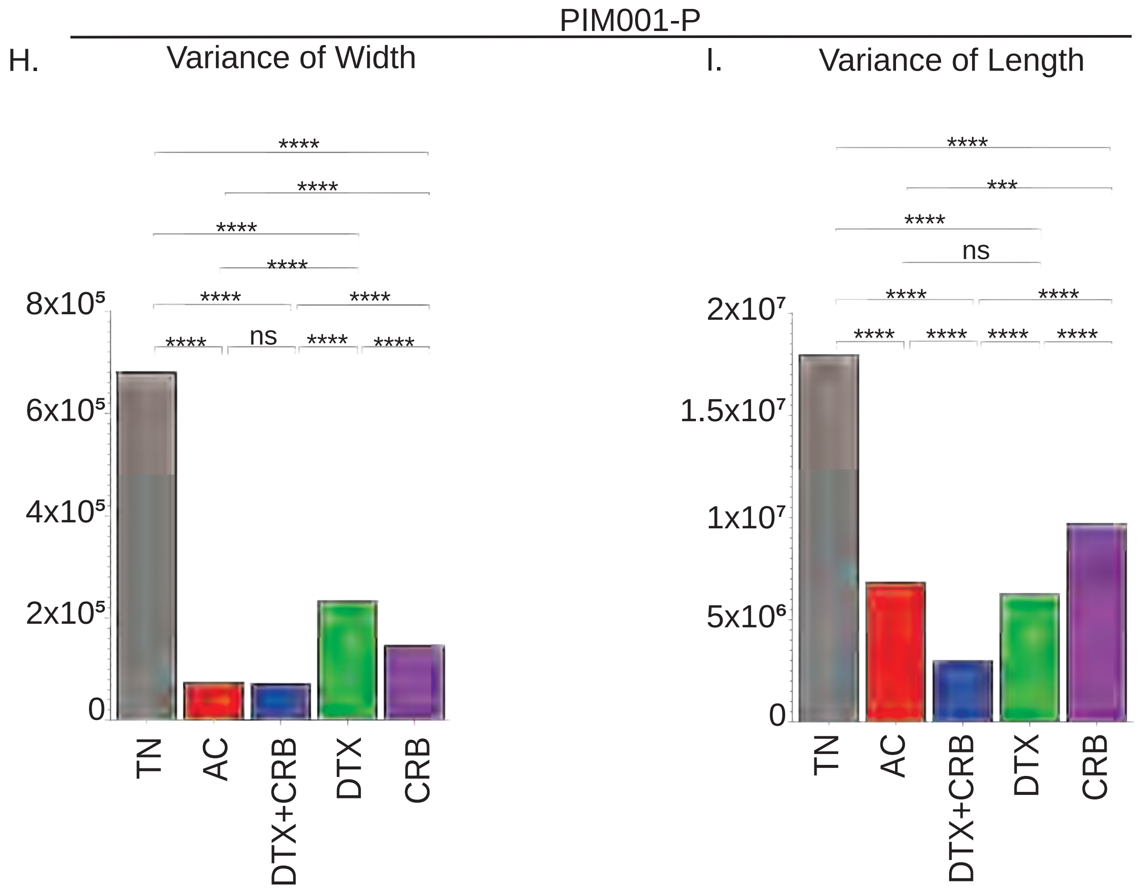


Figure S9. mITH was significantly reduced upon chemotherapy for several mitochondrial metrics in PIM001-P.

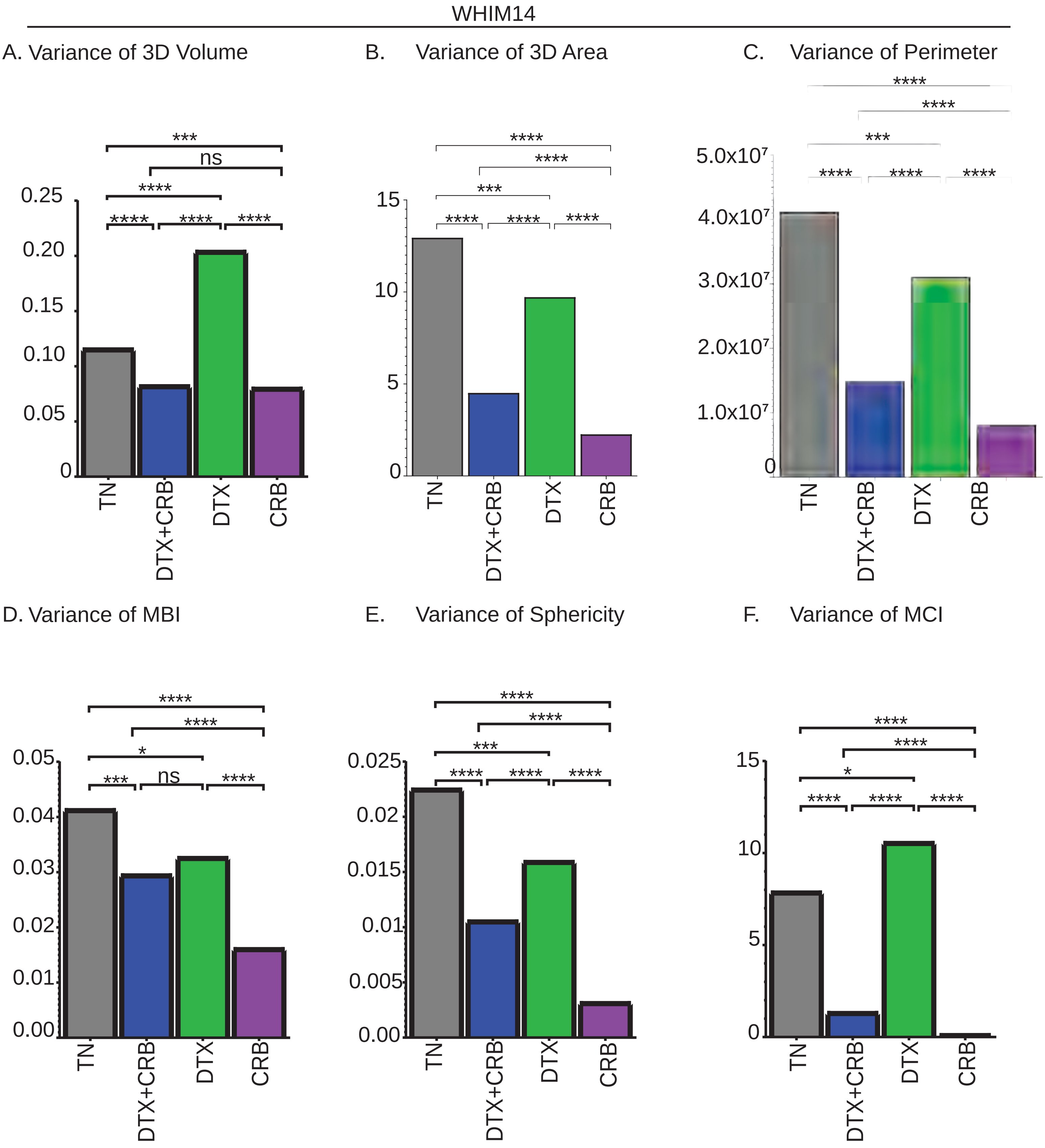


Figure S10. WHIM14 treated with DTX+CRB or CRB only had a significant reduction in mITH.

WHIM14

G.

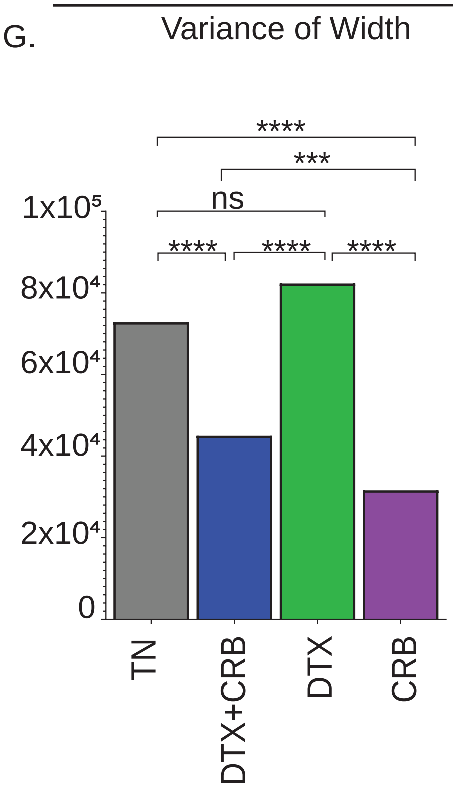


H.

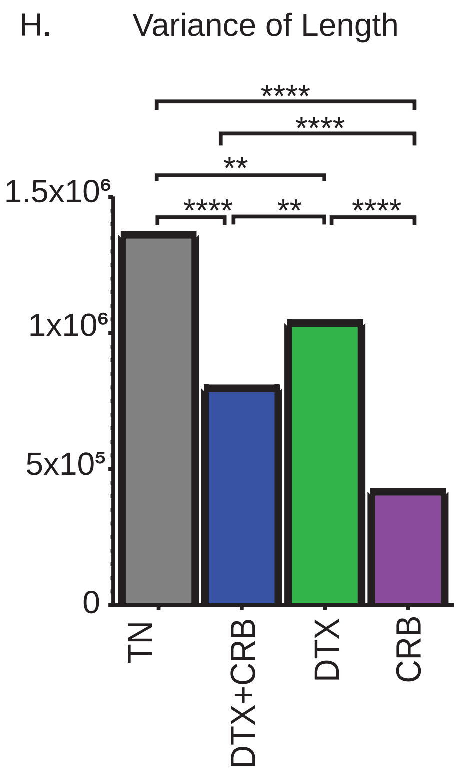


Figure S10. WHIM14 treated with DTX+CRB or CRB only had a significant reduction in mITH.

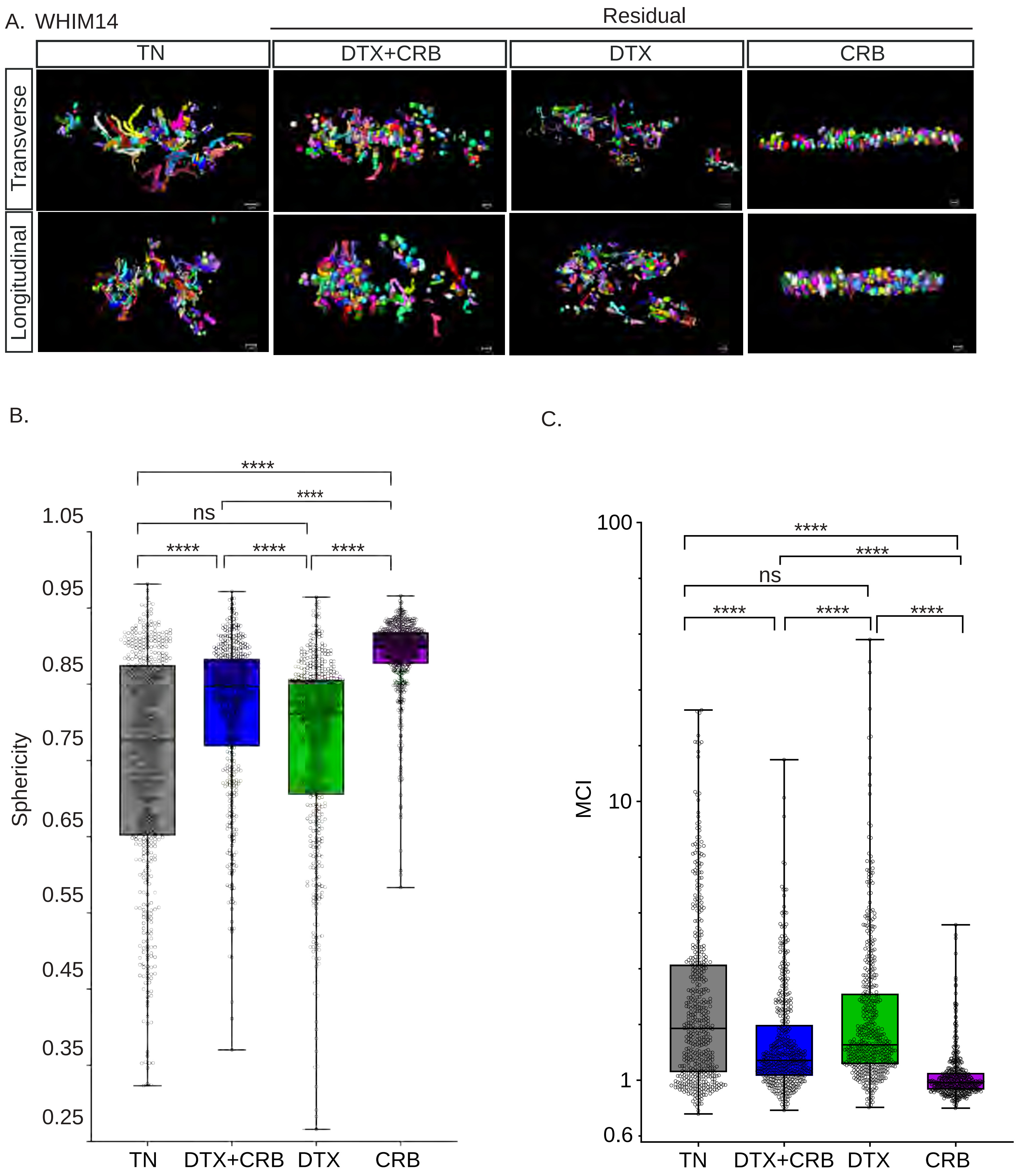


Figure S11. WHIM14 resiudal tumors' mitochondria are less branched and more spherical compared to TN tumors.

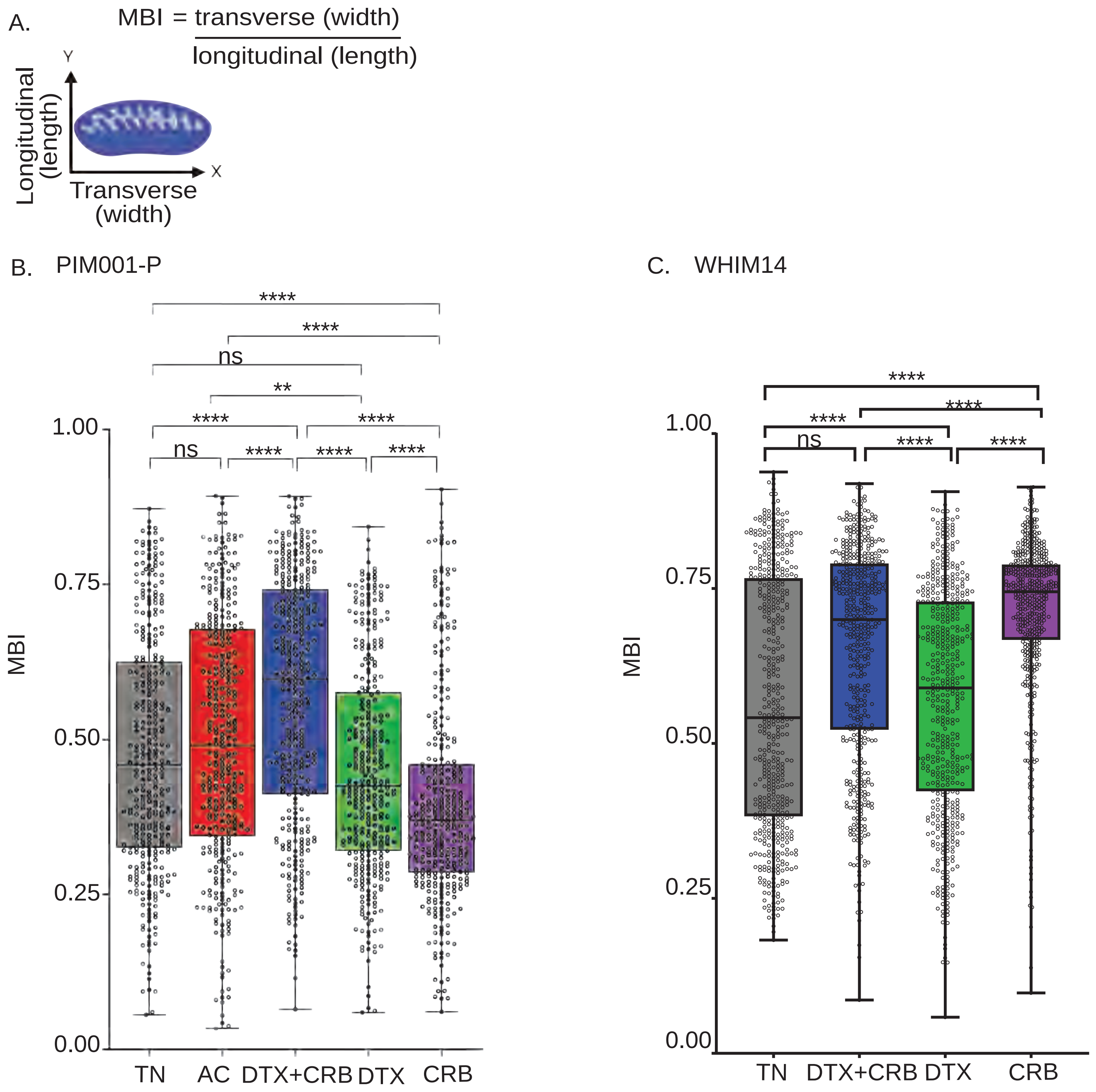


Figure S12. MBI of PIM001-P and WHIM14

A.

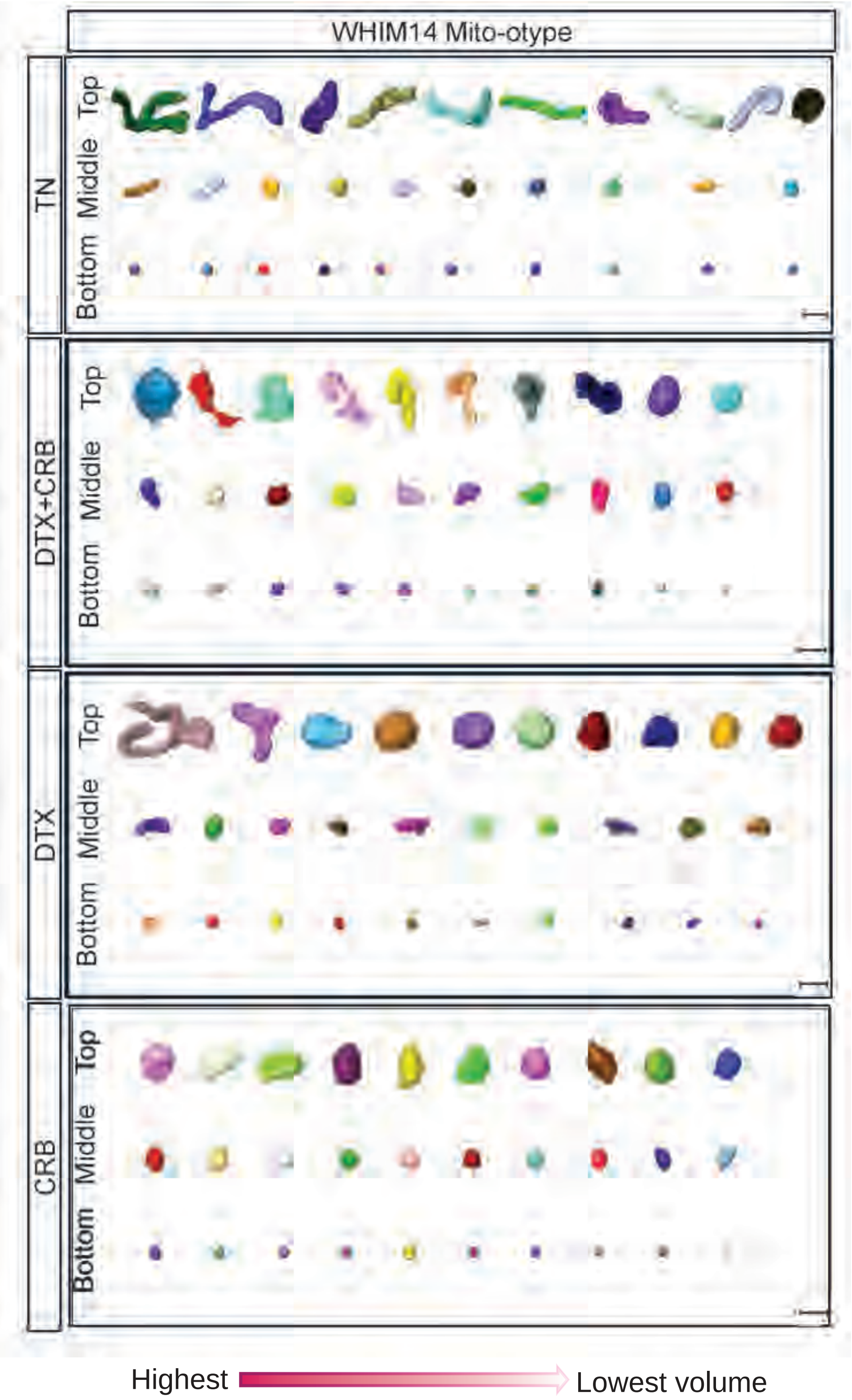


Figure S13. WHIM14 mito-otyping of the diverse mitochondrial structures observed within each treatment group.

WHIM14 lipid droplets

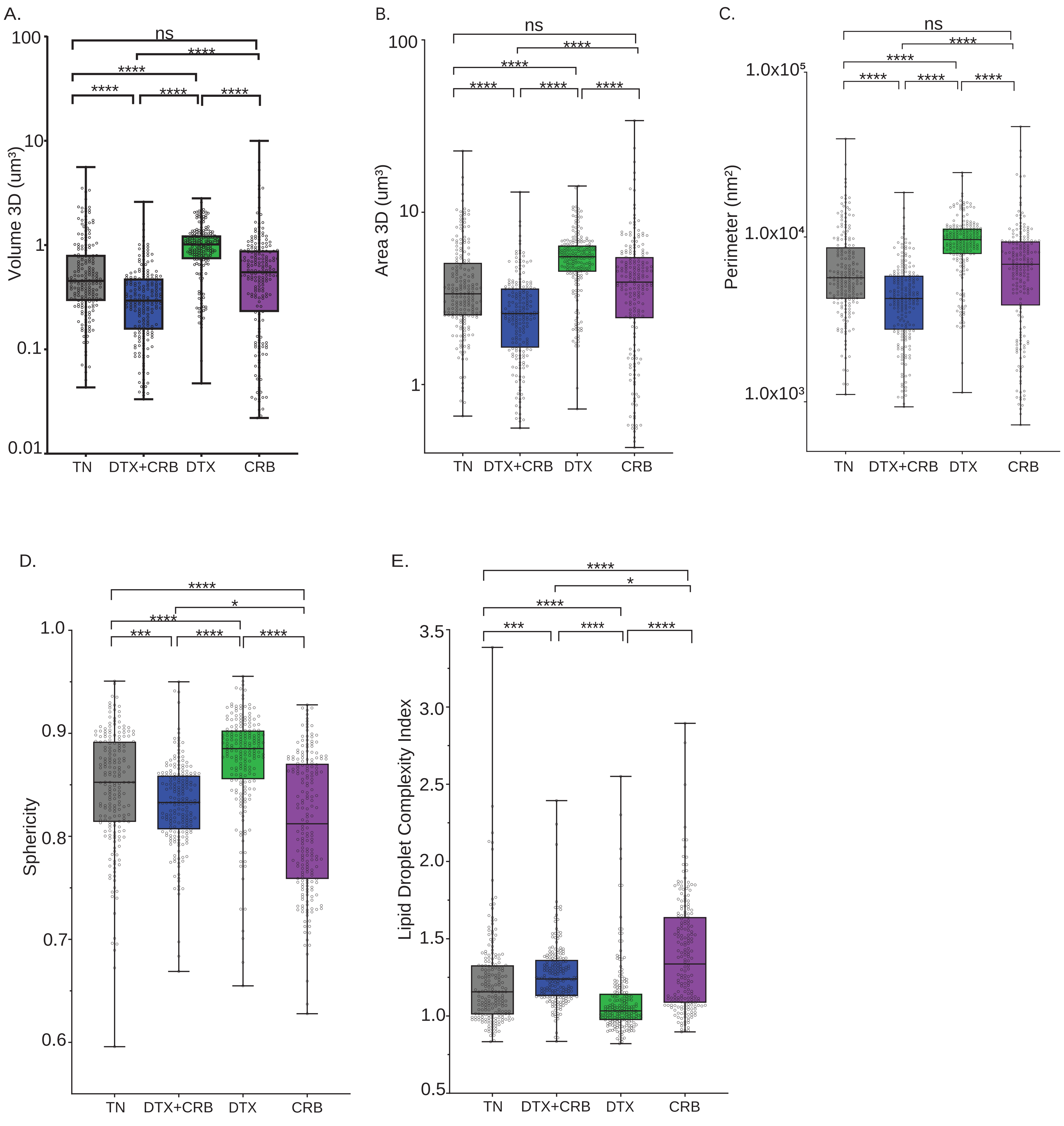


Figure S14. WHIM14 lipid droplet sizes increase after single-agent chemotherapy treatment.
